# Assessing the Generalizability of Machine Learning and Physics-Based Methods with DNA-Encoded Libraries

**DOI:** 10.64898/2026.04.18.719394

**Authors:** Marissa Dolorfino, Daniel Santos Perez, Yao Fu, Shu-Hang Lin, Sean McCarty, Matthew J. O’Meara, Terra Sztain

## Abstract

Predicting protein-ligand binding is a central challenge in computational drug discovery, and while machine learning (ML) and co-folding methods have advanced rapidly, their ability to generalize beyond training or parameterization regimes remains insufficiently understood. DNA-encoded libraries (DELs) enable ultra-large screening of billions of molecules simultaneously, providing a useful testbed for evaluating these approaches at scale. A recent NeurIPS competition revealed that even top performing ML models trained on DEL data failed at generalizing to out-of-distribution (OOD) chemical space. We investigated whether integrating structural modeling could bridge this generalization gap. We systematically assessed state-of-the-art ML, docking, and co-folding methods including Schrodinger Glide, Rosetta GALigandDock, and Boltz-2 with three biologically diverse protein targets screened against libraries containing multiple DEL synthesis formats. While ML excels in-distribution, OOD hit discrimination is dependent on both the target and ligand context, with no single method consistently dominating. These findings demonstrate that benchmark performance alone is insufficient to predict OOD performance, highlighting the need for system-dependent evaluation of binding prediction methods. We provide an open-source package for assessing protein-ligand prediction methods and analyzing high-throughput screening data: DEL-iver.

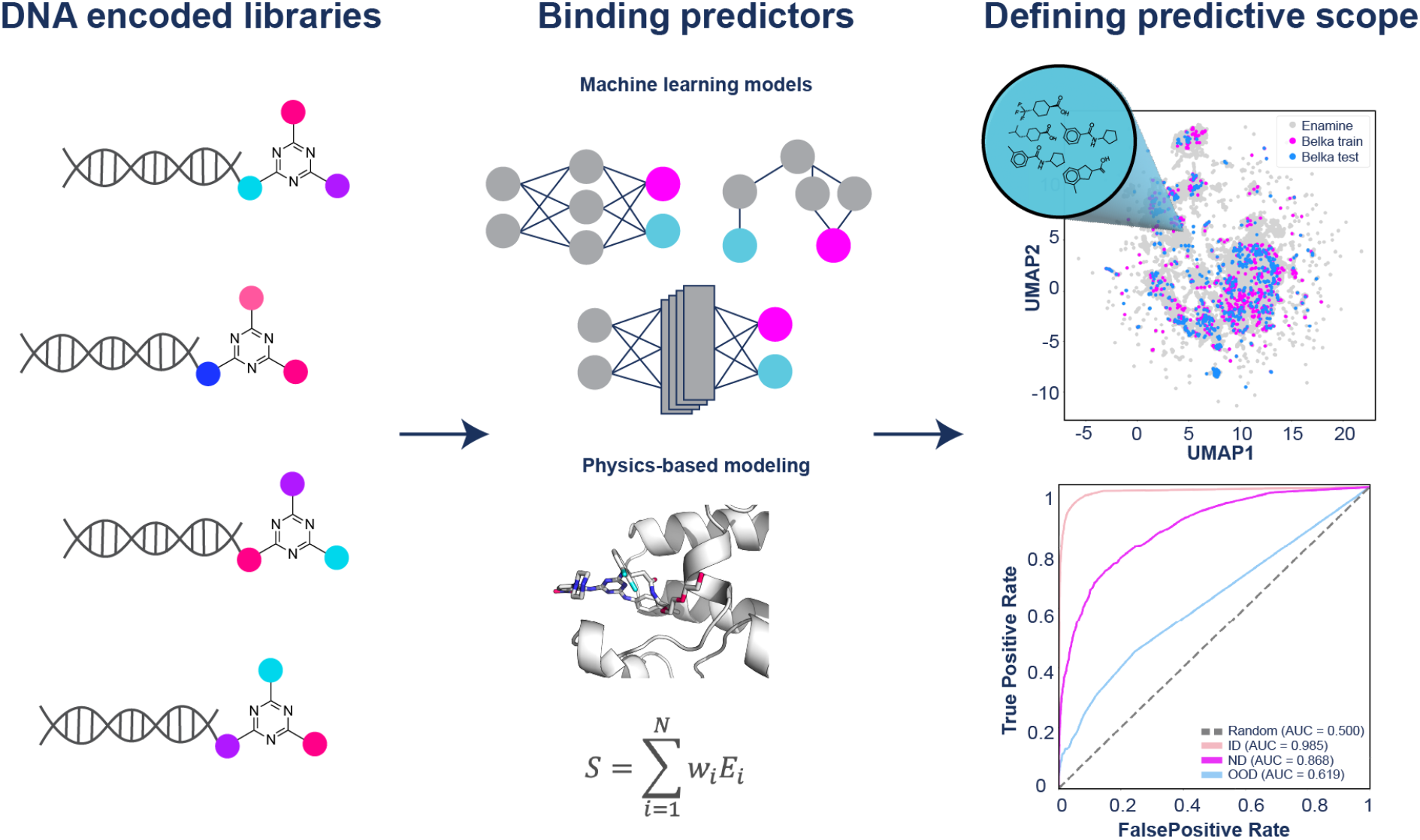

## Introduction

Protein-ligand binding prediction, including the prediction of binding affinity and bound complex structure, plays a central role in computational drug discovery.^1–3^ Traditional approaches include ligand-based QSAR and machine learning (ML) models that infer activity from chemical similarity, as well as structure-based methods such as pharmacophore modeling that identify spatial arrangements of interaction features associated with binding, and molecular docking methods which estimate binding poses and relative affinity using physics-based and empirically derived score functions.^3–5^ Recently, deep learning approaches that jointly model protein-ligand interactions and three-dimensional complex formation have demonstrated highly accurate protein-ligand structure prediction capabilities.^6–10^ Boltz-2, in particular, has extended these advances to predict binding affinity with comparable accuracy to free energy perturbation calculations.^8^

While modern approaches have substantially improved predictive performance, important limitations remain across all method classes. QSAR and ligand-based ML approaches often depend strongly on the composition of the training data and may generalize poorly to novel chemical scaffolds or protein targets.^2,11^ Docking approaches rely on simplified scoring functions and approximations to molecular interactions, which can limit agreement with experimentally observed binding modes and affinities.^12^ Though deep learning methods show great promise, comprehensive benchmarking under diverse and out-of-distribution conditions remains limited.

High throughput screening (HTS) data provide a useful basis for both training predictive models and benchmarking their performance against experimental measurements. Ultra-high throughput screening (uHTS) technologies, such as DNA-encoded libraries (DELs), are an emerging technology for hit-identification, facilitating screening of large, multimillion and billion-compound libraries.^13–18^ DELs are synthesized via combinatorial chemistry coupled with covalent DNA tagging to track the synthetic history of each molecule, enabling simultaneous screening of compounds which can be identified through next-generation DNA sequencing.^17,19^ Key challenges associated with DEL compounds include the complex and expensive off-DNA synthesis of identified hits.^20^ One method to overcome this is to train ML models on DEL screening data, as introduced by McCloskey et al.,^21^ which can then be transferred to predicting diverse, off-DNA purchasable analogues.^21,22^ Reliable implementation of this strategy, however, requires models that can accurately and robustly generalize to unseen molecules. Therefore, rigorous benchmarking efforts to characterize the capabilities and limitations of current methods are needed, especially for this unique compound class which often include larger molecules and occupy distinct regions of chemical space.^23–26^

This has motivated several groups to release public DEL datasets,^22,27^ and sparked the NeurIPS 2024 Kaggle competition - predict new medicines with the Big Encoded Library for Chemical Assessment (BELKA) hosted by Leash Biosciences.^28,29^ The BELKA dataset consists of ∼133 M DEL molecules screened against three targets: soluble epoxide hydrolase (sEH), bromodomain-containing protein 4 (BRD4), and human serum albumin (HSA), resulting in one of the largest experimental affinity selection datasets publicly available. This competition concluded that none of the teams, out of just under 2,000, trained a model that could successfully generalize to the private test set. Using a mean average precision (mAP – the mean of average precision values across each protein group and molecule set) score metric for assessing classification performance, the top scoring model achieved a mAP score of just 0.36 on the private test set, indicating that none of the teams achieved strong out of distribution (OOD) performance, highlighting a limited generalizability of current approaches.

In this work, we leverage DEL uHTS to assess the generalizability of protein-ligand binding prediction methods. First, we summarize the key features of top performing ML approaches from the BELKA generalization competition, noting that a common theme amongst the best scoring models is the use of simple encodings and shallow networks to prevent overfitting. Next, we extend the performance analysis beyond that of the competition by investigating the relationship between data splitting strategies and model accuracy, and compare additional metrics of performance. Critically, we compare several state-of-the-art methods including Rosetta GALigandDock^30,31^, Schrödinger Glide,^32,33^ and Boltz-2^8^ for hit discrimination and enrichment against purely ligand-based ML methods, clarifying the strengths and limitations of each. We also test whether integrating physics-based features in ML model training increases performance accuracy. Finally, we integrate this workflow into an open-source software package for analyzing, modeling, and expanding hits from HTS data, with particular emphasis on the additional challenges posed by DEL screens. This work provides a framework for training machine learning (ML) models on HTS data and integrating molecular docking into hit identification workflows, along with a user-friendly tool for analysis and modeling.

## Results and Discussion

### Insights from Large-Scale Generalization Challenge in Binding Prediction

The BELKA competition provided training data consisting of binary binding labels for a 100M compound DEL screened against three protein targets. Teams were tasked with predicting hit probabilities for molecule–target pairs in a 500k compound test set and were evaluated using mean average precision (mAP) across targets and test splits. Several of the top performing teams provided solutions detailing their successful and unsuccessful strategies. Though one common trend favoring simple encodings and shallow architectures to prevent overfitting emerged, diverse modeling strategies gave similar results reflective of a convergent performance plateau. The results underscore that generalizability remains an unsolved challenge in the field of molecular property prediction, and further advances in modeling strategies and diverse training data are needed.

The first place solution, developed by Victor Shlepov,^34^ was pre-trained in two stages; the first stage consisted of masked language modeling (MLM) while the second used the same model with a separate head to encode Simplified Molecular Input Line System (SMILES)^35^ strings as ECFP4 chemical fingerprints.^36^ Notably, this model was trained on an external dataset^27^ in addition to the BELKA training set, likely aiding in OOD performance. The fifth place solution used an ensemble of three net architectures: CNN, transformer, and mamba state space model.^37^ The input was treated as a sequence, and the task was a 3-class (BRD4, HSA, sEH) multi-label problem.^38^ The eighth place solution used a multi-target transformer that utilized pretrained models MolFormer and RoBERTa.^39,40^ The model was trained independently on shared and non-shared building block splits.^41^ A comprehensive description of publicly available solutions can be found on the challenge website.^29^

Our team tested various molecular encodings including ECFP4,^42^ MACCS,^43^ and APDP chemical fingerprints,^42^ molecular graphs, and one-hot encoded SMILES strings, and ML architectures including random forests, graph convolutional networks (GCNs), and multi-layer perceptrons (MLPs). A detailed summary of each model is provided in the Methods and **Table S1**. Two unique features of our models which noticeably improved performance include 1) concatenating fingerprint representations ECFP4 and APDP and 2) sharing layers between building blocks two and three to encode permutation invariance accounting for rotational freedom around the DNA connected building block one. We did not find evidence for these strategies being used by others in the competition.

Our objective in the following sections is to isolate and evaluate the impact of modeling choices such as data splitting, augmentation, downsampling, and physics-based methods. Therefore, we standardized the model architecture and training procedure across experiments rather than performing extensive hyperparameter optimizations. Unless otherwise noted, subsequent models described used this fixed MLP architecture (see Methods).

### Generalizability Across Data Distributions and Targets

Since the BELKA competition prioritized generalizability, a test set was designed containing building blocks (BBs) and scaffolds unseen in the training set. These unseen molecules can be grouped into one of three categories: 1) molecules composed of unique combinations of the BBs seen in the training surrounding a triazine core scaffold (in distribution, ID), 2) molecules composed of BBs unseen in the training surrounding a triazine core scaffold (near distribution, ND), and 3) molecules composed of BBs unseen in the training without a triazine core (out of distribution, OOD). Of the total molecules in the test set, 66% belong to the ID group, 4% the ND group, and 30% OOD (**Figure 2A**). The average Tanimoto similarity is lower for groups further from the training distribution (**Figure 2B**), and only compounds in the ID group show an overlap of ECFP4 fingerprints plotted in UMAP space (**Figure 2C**). As expected, the model performance drops for groups with increasing divergence from the training set (**Figure 2D-I**). This performance, however, varies for each protein and greater performance on the ID set does not always correlate with greater performance on the ND or OOD sets.

**Figure 1.**
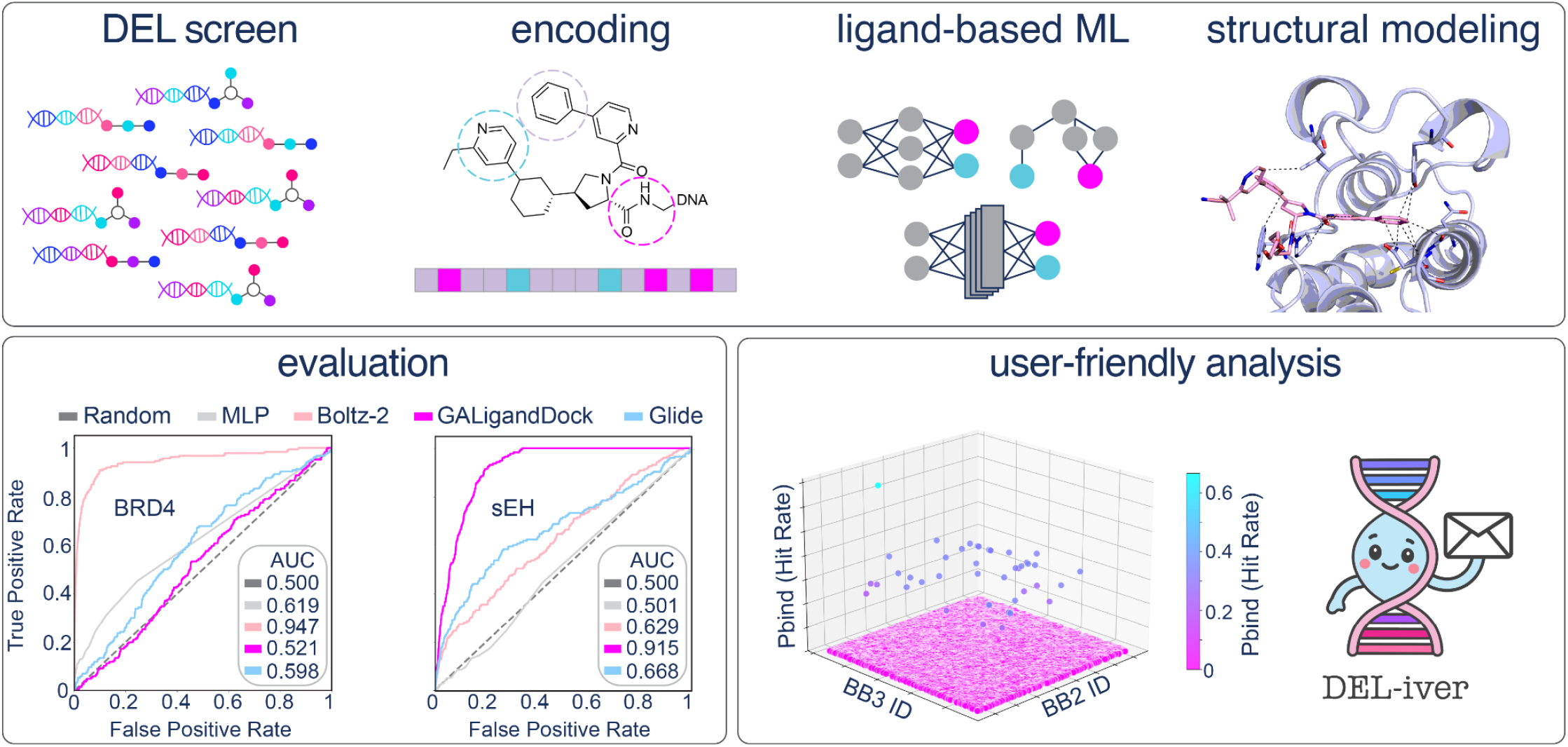
Schematic overview of this study. Representations showing DEL screening compounds, molecular encodings, ligand-based ML, structural modeling, evaluation, and our user-friendly analysis tool, DEL-iver.

**Figure 2:**
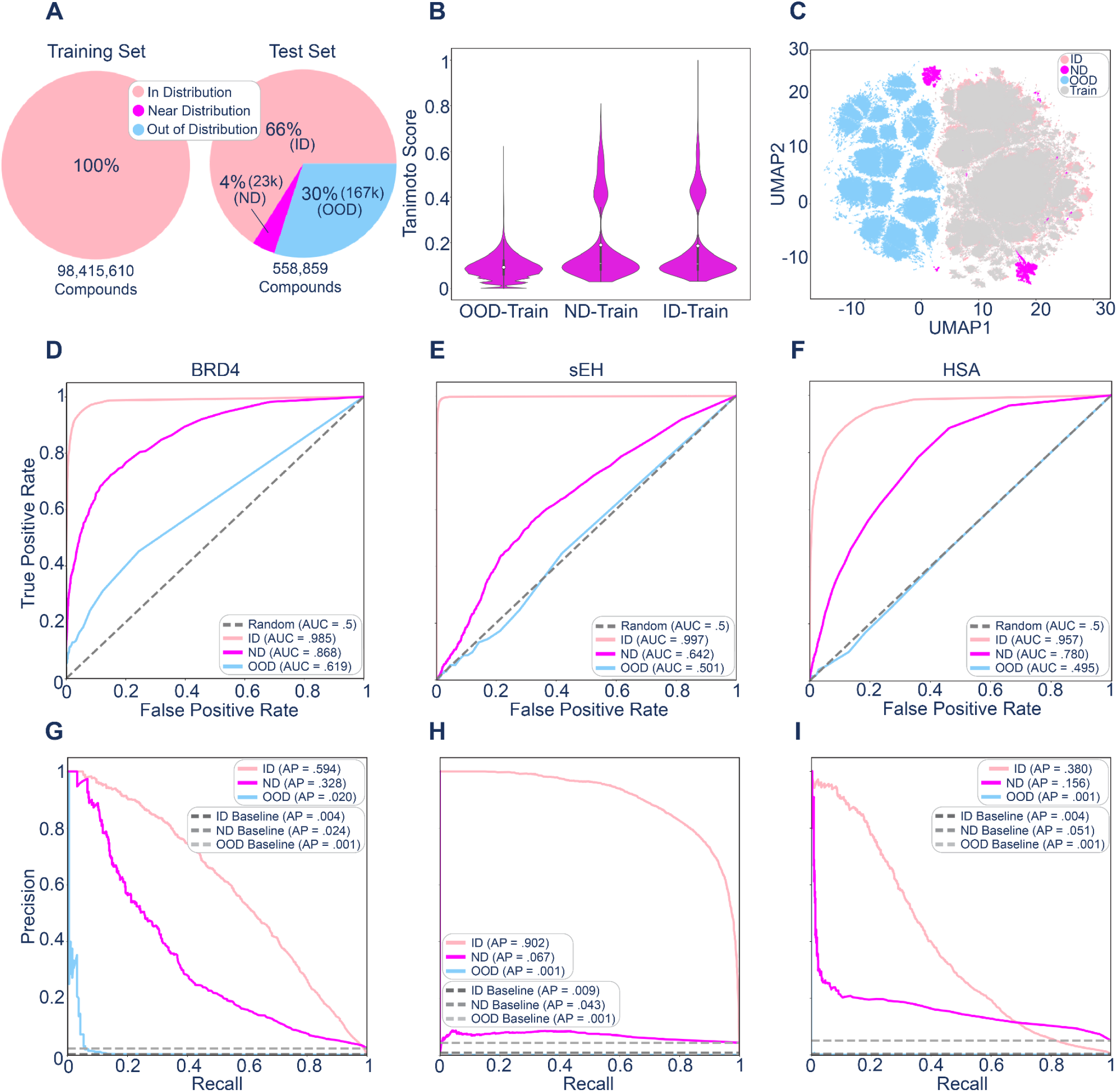
Composition of BELKA dataset and baseline ML performance BELKA dataset. **A)** Distribution of In Distribution (shared scaffold, shared building blocks; ID), Near Distribution (different scaffold, shared building block; ND), and Out of Distribution (different scaffold, different building blocks; OOD) sets in the BELKA train and test sets. **B)** Tanimoto similarity of ECFP4 fingerprints between molecules in the ID, ND, and OOD sets and the train set. **C)** Uniform Manifold Approximation and Projection (UMAP)^44^ of the ECFP4 chemical fingerprints of the ID, ND, and OOD molecule sets. **D-F)** Receiver Operating Characteristic plots comparing performance of our baseline ML model on the BELKA test set stratified by ID, ND, and OOD for BRD4, sEH, and HSA, respectively. **G-I)** Precision-Recall plots corresponding to the same models and test sets shown in D-F. Baseline corresponds to the number of positive hits out of the total data in each test set.

For example, predicting binding of ID molecules to sEH gave the highest average precision (AP) of 0.902, followed by 0.593 for BRD4, and 0.380 for HSA, with corresponding area under the receiver operating characteristic curve (AUROC) values of 0.997, 0.985, and 0.957, respectively (**Figure 2D-I**). For prediction of ND molecules, the mAP values for sEH, BRD4, and HSA were 0.067, 0.328, and 0.156, and the AUROC values were 0.642, 0.868, and 0.780, respectively (**Figure 2D-I**). The relatively low AP of sEH on this set, coupled with such high ID performance indicates greater overfitting to the ID set than the BRD4 and HSA models. This may be attributed to the higher hit rate of sEH in the test set (0.85%) compared to the hit rates in the test set for BRD4 and HSA of 0.40% and 0.50%, respectively, or differences in data quality, including signal to noise ratio. For prediction of OOD molecules, the AP was 0.020 for the BRD4 model, and 0.001 for both sEH and HSA, while AUROC values for sEH, BRD4, and HSA were 0.501, 0.619, and 0.495, respectively (**Figure 2D-I**). Taken together, all models showed greatest performance on the ID sets and near random performance on OOD.

As an additional test, we sought to evaluate an independent set of OOD molecules not connected with the competition. A published DEL screen against sEH was previously conducted by Zhang et. al,^27^ using a DEL constructed by attaching three building blocks linearly, rather than around a central scaffold. We evaluated the sEH model on this dataset, and saw only slightly higher performance than on the OOD set from the competition, with an AP of 0.017 and AUROC of 0.545 (**Figure S1**). This demonstrates that the OOD set was not uniquely challenging, but the model performs poorly at generalization far from the training distribution.

### Effect of Modifying Dataset Compositions

We sought to further investigate the role of data composition on model performance. One of the obvious imbalances in DEL screening data is the large proportion of non-hit molecules compared to hit molecules. In the BELKA training set, hits make up only 0.54% of the total molecules. We tested the impact of downsampling the overrepresented non-hit molecules by randomly removing varying proportions from training, and evaluating models on the original BELKA test set. Surprisingly, a drop-off in performance was not detected until 99% of the non-hit data was removed, decreasing the training data from ∼100M to ∼1M datapoints, with a new hit composition of 54.14% (**Figure 3A**). This suggests that dataset size alone is not the primary determinant of model performance, but the composition and quality of such datasets are also critical. The significant decrease in training set size, however, opens up the possibility of much faster training, enabling more complex, detailed chemical representations and architectures to be explored.

**Figure 3:**
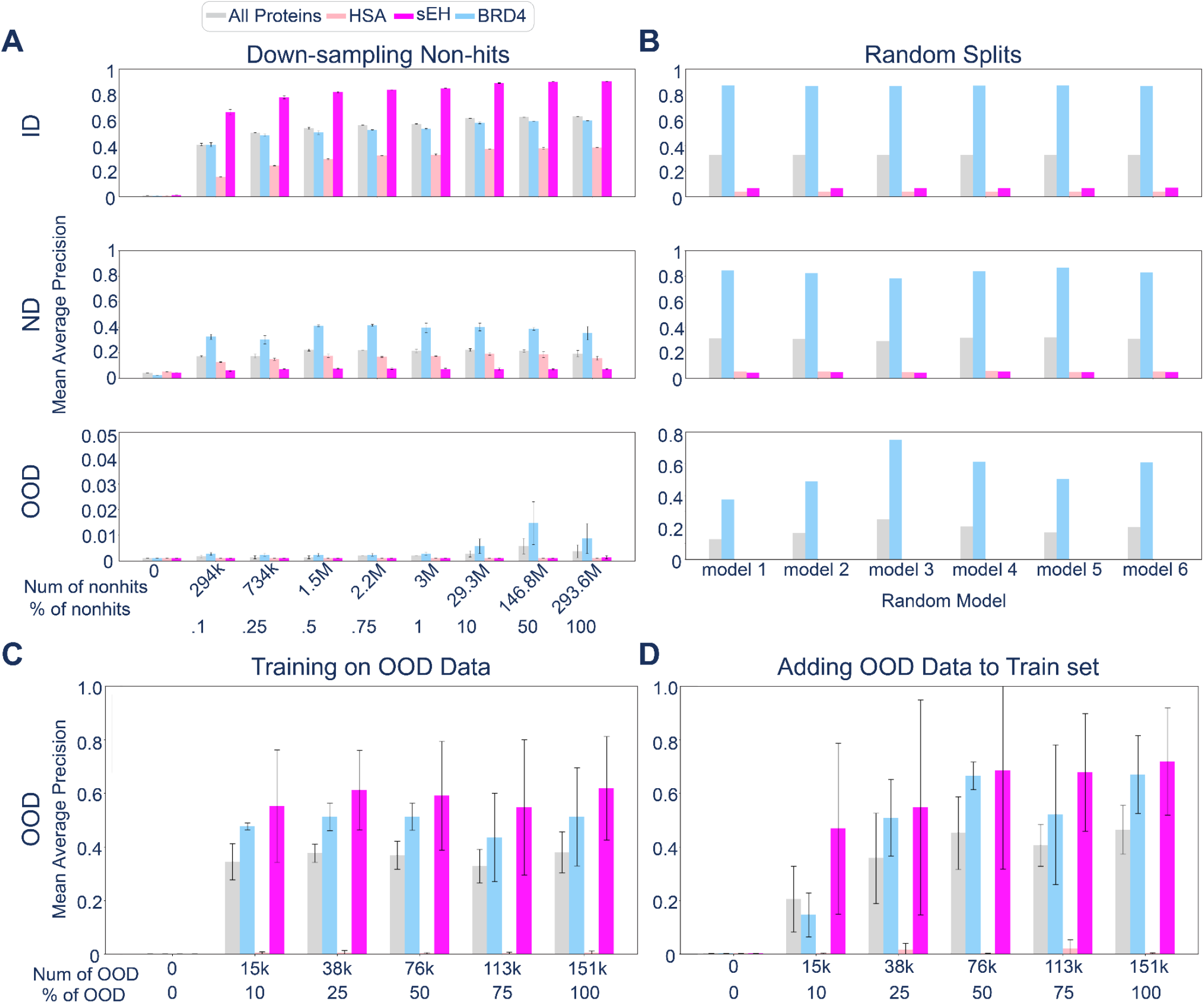
Performance of baseline ML model with modified train and test sets. **A)** Mean average precision of baseline model trained with decreasing amounts of non-hit molecules from the original BELKA training set. Error bars represent averages and standard deviation across three training replicates. Performance is shown for the ID, ND, and OOD sets. **B)** Performance of baseline ML model trained on random splits of BELKA data with 90% non-hits removed and 20% kin0 maintained in the test set. Six independent models were trained and performance is shown on each random test set with ID, ND, and OOD labeled based on classification in the original BELKA train/test split. **C)** Performance of baseline ML model on a constant withheld kin0 test set (∼15k molecules) trained on random subsets of the remaining kin0 molecules. Three independent models were trained with independent random subsamples for each of the percentages shown. Error bars show the average and standard deviation across the three models. **D)** Performance of baseline ML model on the same withheld kin0 test set as in **C** trained on subsets of kin0 molecules aggregated with the original BELKA train set. The performance is averaged across the three models, and standard deviation bars are shown.

We next compared performance on random train-test splits of the aggregated BELKA train and test data. Random splits are commonly used in ML,^45–47^ and they can also help test whether diversification improves performance on the original OOD set of non-triazine containing molecules. We refer to the non-triazine group as “kin0” in this section since they will be seen during training and are therefore no longer OOD. We trained six models with 80/20 random train/test splits of the aggregated BELKA dataset. Non-hits were downsampled, removing 90% from the original training set, before aggregation. To ensure even distribution of kin0 and non-kin0 molecules, we ensured the random split test set contained 20% of the kin0 molecules. As expected, for BRD4, the presence of kin0 compounds during training improved predictions on the held-out kin0 molecules, with APs increasing from 0.020 to a mean of 0.558 across the six random models and AUROC increasing from 0.619 to a mean of 0.968. Interestingly, performance also improved for non-kin0 molecules in the BRD4 model, with AP values increasing from 0.594 to 0.870 on average, and AUROC from 0.985 to 0.989 on average across the six models for ID, and AP from 0.328 to 0.831 on average and AUROC from 0.868 to 0.989 on average for ND (**Figure 3B**). This suggests that the dataset diversity may lead to more generalizable representations that better capture the underlying structure of the data.

For sEH and HSA however, the random splits did not improve performance on kin0 molecules, and the performance on other molecule sets substantially decreased. For sEH, AP decreased from 0.902 to a mean of 0.067 for ID and from 0.067 to a mean of 0.045 for ND. AUROC also decreased from 0.997 to 0.499 on average for ID and 0.642 to 0.508 on average for ND. The HSA AP decreased from 0.380 to a mean of 0.039 for ID and from 0.156 to a mean of 0.050 for ND, and AUROC decreased from 0.957 and 0.780 for ID and ND, to 0.500 and 0.501 on average, respectively (**Figure 3B**). In this case, the training set diversity degraded model performance, possibly indicating a greater sensitivity to distributional broadening or noise than the BRD4 model.

With random splitting, the kin0 compounds make up less than 2% of the total training data, therefore we investigated whether training only on kin0 compounds could maximize predictions in this region of chemical space. We evaluated performance on a constant 15k kin0 compound holdout set, while training on increasing amounts of the remaining kin0 data from 15k to 151k. The training set size minimally impacted the AP, with values near 0.6 for sEH, 0.4 for BRD4, and 0.001 for HSA regardless of training set size (**Figure 3C**). The AUROC was also minimally impacted by training set size, with values near 0.99 for BRD4 and sEH, and values near 0.55 for HSA. This large performance gap, in which models learn to predict kin0 binders to BRD4 and sEH but not to HSA may be due to variance in noise for each target, and underscores that similarity to the training distribution alone does not guarantee accurate predictions.

Next, the original BELKA training molecules were aggregated with increasing amounts of kin0 training data to test whether significantly increasing the training set size could improve performance in this lower data regime, a phenomenon we observed previously with deep mutational scanning data.^48^ While performance with sEH and BRD4 apparently improves with increasing amounts of kin0 data, large variance across training replicas prevents a definitive conclusion (**Figure 3D**).

### Structural Modeling Improves Out-of-Distribution Hit Discrimination

To examine the impact of structural modeling on hit discrimination performance, we compared state-of-the-art docking software including Schrödinger Glide,^32,33^ Rosetta GALigandDock,^49,50^ and the co-folding foundation model Boltz-2^8^ to the ligand-based ML predictors. To mitigate excessive computational expense, docking with Glide, GALigandDock, and Boltz-2 was only carried out on BRD4 and sEH with the ∼165k OOD molecules.

Both Glide and GALigandDock require definition of an initial starting conformation or docking grid. Details of structure preparation are outlined in the Methods section. Of note, we included a triethylene glycol (PEG3) linker at the DNA attachment site on BB1 to prevent burial of that building block, which would be sterically prohibited under the experimental conditions we were aiming to predict. Additionally, since sEH contains two domains as possible binding sites, we used Diffdock-L to predict which site each molecule was predicted to bind. Only a very small fraction, 598 molecules, bound to the ATP binding pocket while the majority bound to the substrate pocket.

Performance of each method was evaluated and compared to the baseline ML model on the OOD set. Raw docking scores from GALigandDock and Glide were converted to binding probabilities using a 0 to 1 min-max scaling. This was not necessary for Boltz-2 which directly predicts a 0 to 1 probability of binding. Visually, the probability distribution of hits and non-hits shows near complete overlap between hits and non-hits in most cases, however for BRD4 using Boltz-2, and for sEH using GALigandDock distinct peak centers were observed with hits around 0.8 and 0.9, respectively, and non-hits around 0.2 and 0.3, respectively. In the case of BRD4 with Boltz-2, the distributions are partially overlapping, (**Figure 4A**) whereas for sEH with GALigandDock, all hits scored in the high probability region, while non-hits were distributed across both corresponding to a high true positive rate with no false negatives, but several false positives (**Figure 4C**).

**Figure 4.**
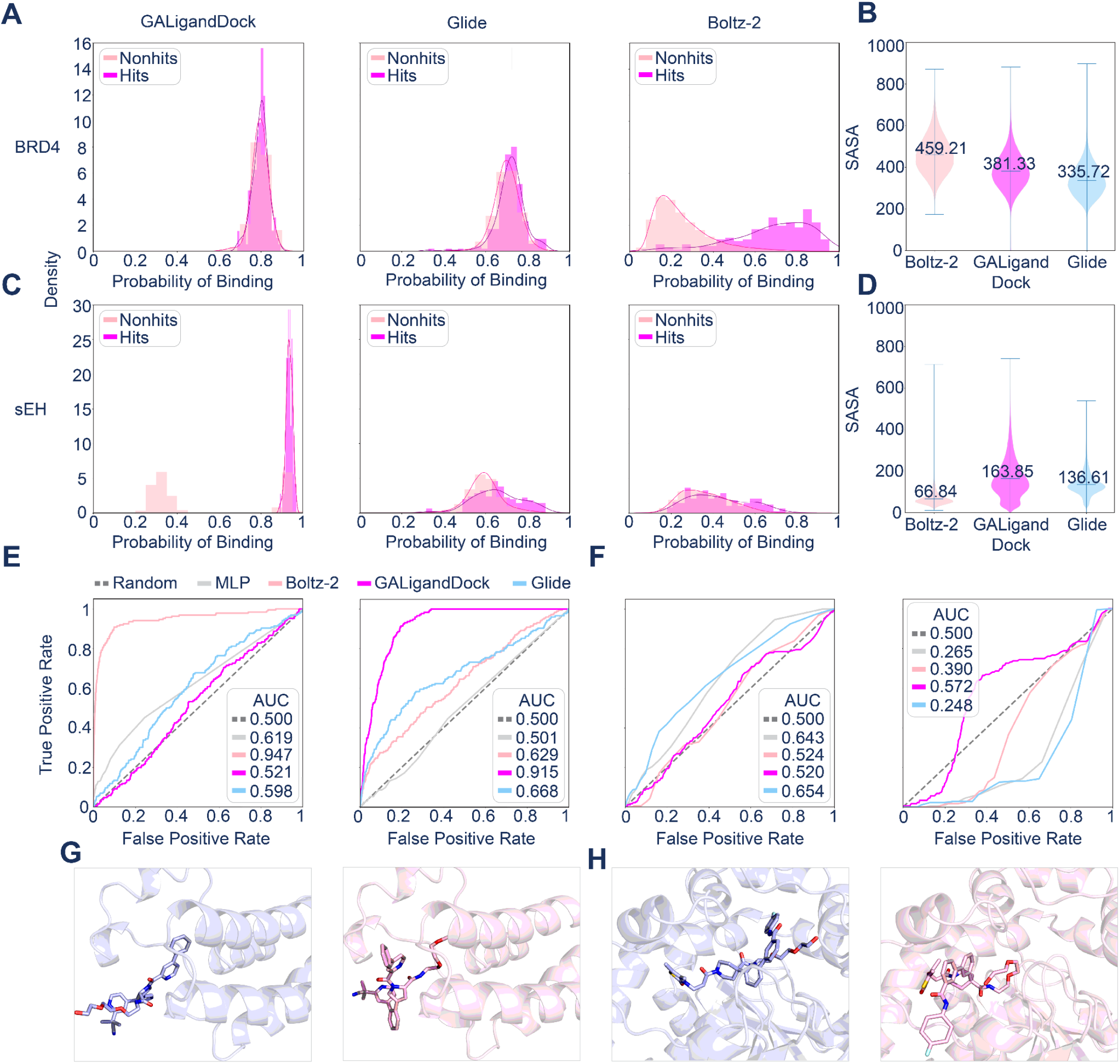
Performance of structure-based methods on the OOD BELKA set and analysis of and training on physics-derived features. **A)** Probability of binding derived from GALigandDock, Glide, and Boltz-2 for the OOD BRD4 set. **B)** SASA of the docked structures from GALigandDock, Glide, and Boltz-2 for the OOD BRD4 set. **C)** Probability of binding derived from GALigandDock, Glide, and Boltz-2 for the OOD sEH set. **D)** SASA of the docked structures from GALigandDock, Glide, and Boltz-2 for the OOD sEH set. **E)** AUROC plots showing the performance of the baseline ML predictions and structure-based methods on predicting hits and non-hits for the BRD4 (left) and sEH (right) OOD sets. **F)** AUROC plots showing the performance of the baseline ML architecture and ML architectures using concatenated PLIP and ECFP4 fingerprints as input. All models here were trained on the 25k ID set and tested on the 165k OOD set (BRD4: left, sEH: right). For each protein, the PLIPs calculated from each structural method were used to train a separate model. **G)** Docked structures from Boltz-2 (purple) and GALigandDock (pink) of an OOD molecule to BRD4 for which Boltz-2 correctly classified the molecule as a non-hit, while GALigandDock incorrectly classified the molecule as a hit. **H)** Docked structures from Boltz-2 (purple) and GALigandDock (pink) of an OOD molecule to sEH for which Boltz-2 incorrectly classified the molecule as a hit, while GALigandDock correctly classified the molecule as a non-hit.

Superior hit discrimination performance of Boltz-2 with BRD4 is reflected by the AUROC of 0.947 compared to 0.619, 0.521 and 0.598 for the MLP, GALigandDock, and Glide, respectively (**Figure 4E**). However, the AP was not improved, with values of 0.001, 0.001, 0.002, and 0.007 for Boltz-2, MLP, GALigandDock, and Glide, respectively, indicating that while Boltz-2 significantly improved hit-discrimination of these compounds with BRD4, the hit-enrichment remained unchanged. Incremental enrichment gains were provided with GALigandDock, and Glide compared to the MLP, with worse overall hit-discrimination.

The superior hit discrimination performance of GALigandDock with sEH is reflected by the AUROC of 0.915 compared to 0.501, 0.629, and 0.668 for the MLP, Boltz-2, and Glide, respectively and AP of 0.009 with GALigandDock and 0.001, 0.003, and 0.003 for the MLP, Boltz-2, and Glide, respectively (**Figure 4E**). Though GALigandDock outperformed other methods for sEH, all structural methods outperformed the baseline MLP.

These results show that docking and cofolding can be used to classify hits with greater out of distribution performance than the ligand only ML models in many cases, however the performance across methods depends on the target. To test whether the optimal target/method pair remains consistent across chemical space, we selected 25k random molecules from each of the BRD4 and sEH ID sets for comparison, including all 1480 BRD4 and 3564 sEH hit compounds. Binding probabilities obtained from docking to this compound set did show distinct peak centers, with the distribution of hit and non-hit binding probabilities mostly overlapping, and there was less of a clear standout method (**Figure S3A**). Boltz-2 remained the optimal method for BRD4 prediction with AUROC of 0.697 compared to 0.469 and 0.525 for GALigandDock and Glide, respectively. AP on this set was also highest for Boltz-2, with an AP of 0.146, compared to 0.056 and 0.064 for GALigandDock and Glide, respectively. For sEH, Glide gave the optimal hit-discrimination performance with AUROC of 0.614 compared to 0.583 and 0.512 for Boltz-2 and GALigandDock, respectively (**Figure S3B**). AP on this set was highest for Boltz-2, with AP of 0.075, followed by APs of 0.064 and 0.061 for GALigandDock and Glide, respectively.

Enrichment factors reflect the ability of molecular docking methods to detect true positives from the dataset compared to random selection at different threshold values for selecting positives.^51,52^ Enrichment factors of each method calculated for the top 0.5, 1, 5, 10, and 20% threshold values for selecting hits can be found in **Table S2**. The largest EF 0.5%, with a value of 86.04 was from Boltz-2 with BRD4 and the OOD ligands set. The next highest EF 0.5% was nearly an order of magnitude lower, with a value of 9.15 for sEH with the OOD ligand set using either GaLigandDock or Glide. Consistent with the other metrics calculated, none of the methods performed well on the ID set, with Boltz-2 and BRD4 giving the highest EF 0.5% value of 4.37, and all others giving values near 1 (**Table S2)**.

Next, we investigated the structural poses generated from each method. Since no ground truth structures exist with these molecules bound to these targets, a reasonable assumption might be that the method with the greater capacity for hit discrimination would produce the more reliable pose prediction. We make these generated structures available at https://huggingface.co/datasets/SztainLab/BELKA_docked_structures for future benchmarking and structure-based drug design efforts. To quantify deviation of predicted ligand poses, pairwise RMSD of the ligand was calculated after receptor alignment across each method. A range of 1-16 Å for BRD4 hits and 1-20 Å for sEH hits is observed for all methods (**Figure S4**). Non-hits showed greater deviations with most ligands ranging between 1-26 Å, though some as high as 70 Å for both BRD4 and sEH when comparing against Boltz-2, where a predefined binding site is not provided. The average ligand solvent accessible surface area (SASA) was largest for BRD4 and Boltz-2 for both the OOD and ID sets, which correlated with the optimal method corresponding to larger SASA (**Figure 4B**). For sEH, GaLigandDock resulted in larger SASAs on average for both sets, while GALigandDock was only optimal for the OOD set, and Glide was superior for the ID set (**Figure 4D**).

To determine whether features derived from structural modeling could improve ML performance, we calculated Protein Ligand Interaction Profiler (PLIP) fingerprints,^53^ which encode the interacting residues for each ligand. As expected, the number of total interactions is inversely proportional to the ligand SASA (**Figure 4B,D, Figure S2**). These physics-informed features were then concatenated with ECFP4 fingerprints to determine the impact on performance of the baseline MLP model. Though there is a slight improvement over baseline with some models, others negatively impact performance, and the results are not consistent with score metrics. For example, the AUROC increased from 0.643 to 0.654 for BRD4 using PLIPs generated from Glide structures, while the AUROC decreased to 0.524 with PLIPs obtained from Boltz-2, despite Boltz-2 binding probabilities giving greater hit discrimination for BRD4. For sEH, PLIPs generated from GALigandDock showed the greatest improvement over baseline, with AUROC increasing from 0.265 to 0.572, consistent with GALigandDock outperforming other methods at hit discrimination for sEH (**Figure 4F, Figure S3**). These results suggest that the method most capable of hit discrimination does not necessarily lead to the most optimal features for ML.

### Open-Source DEL-iver Analysis and ML Package

To facilitate reproducibility, future developments, and user-friendly analysis of HTS, particularly DEL screening data, we compiled code used in this study into an open-source python package, DEL-iver. In addition to conducting the above research, preparing screening data for ML, training and evaluating models, we have included features we hope will be useful for a range of users from non-computational chemists to ML experts (**Figure 5**).

**Figure 5.**
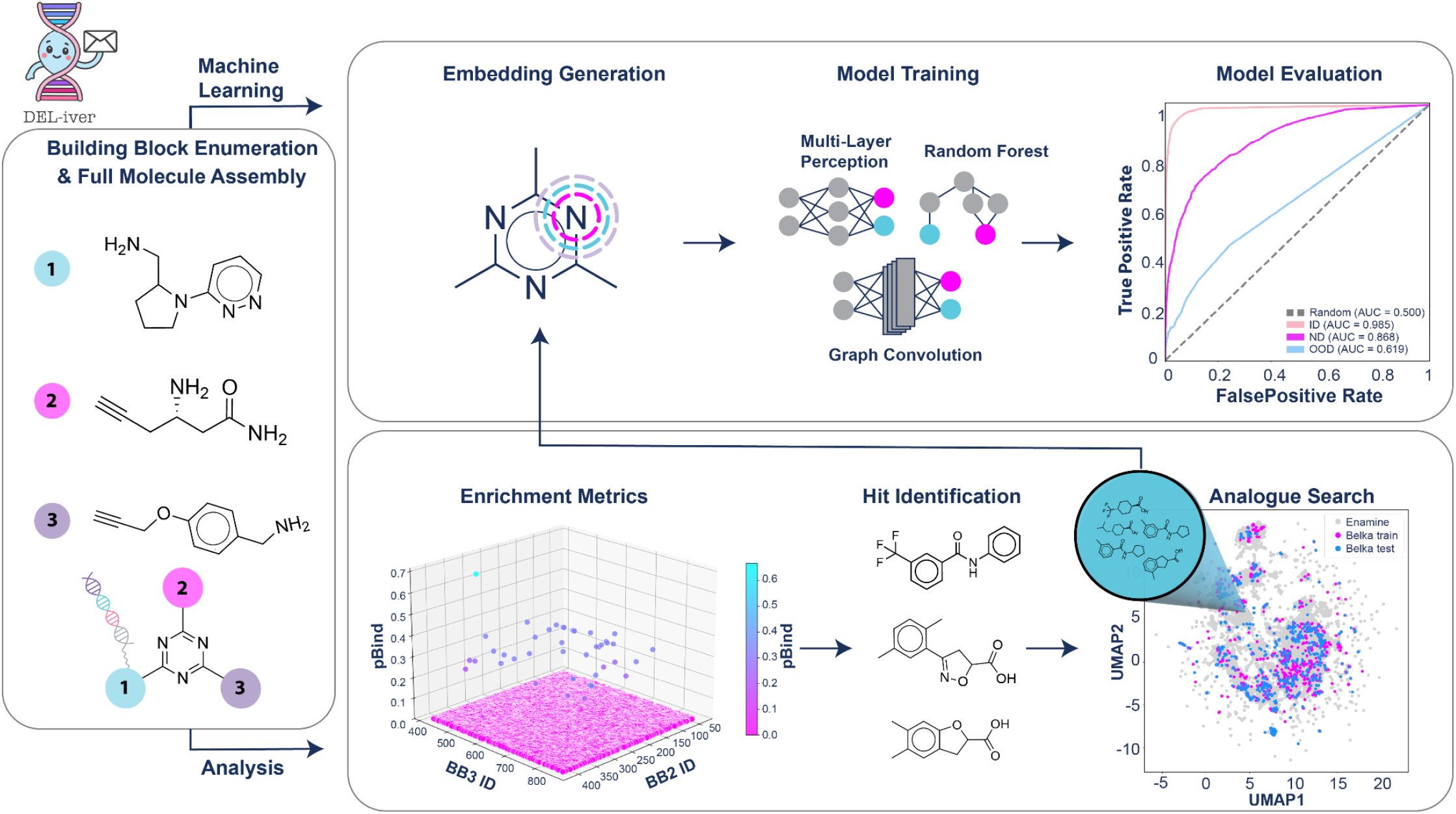
Overview of the DEL-iver package. DEL-iver enables streamlined analysis of DNA-encoded library (DEL) datasets. Building blocks are first assigned unique numerical identifiers, and full molecules can optionally be assembled on a triazine core scaffold. The workflow supports flexible usage, including standalone analysis, standalone machine learning (ML), or an end-to-end analysis-to-ML pipeline. The analysis module computes and visualizes enrichment metrics, including pBind and enrichment, and facilitates identification and rendering of top-ranking molecules. These hits can be used for analogue searching within external chemical databases. The resulting compounds can then be directly integrated into the ML pipeline, where molecular embeddings are generated and diverse models are trained, all with built-in model evaluation.

To demonstrate the functionalities of DEL-iver, we further analyzed the BELKA dataset, revealing distinct, target-specific structural patterns among hit compounds. As the DNA-bearing moiety, BB1 was excluded from analysis.^27,54^ Analysis was carried out at the level of building blocks and disynthons. Each was filtered to include only those observed at least 30 times per protein target.^27^ From BB2 and BB3 and pBind was calculated for building blocks (**Figure S5**) and disynthons (**Figure S6**). The *Del_iver_results*.*py* script takes in raw csv files to generate these graphs, provides enrichment statistics (**Table S3**), and outputs the top 10 chemical structures (**Figure S7-9**). Visual inspection of these structures revealed that among the top sEH candidates, three featured spirocyclic rings and eight contained aromatic heterocycles (**Figure S7)**. The top 10 performing disynthons for BRD4 were characterized by 5 being sulfur-containing heterocycles, 3 containing aromatic halides and 9 were nitrogen-bearing bicyclic or fused-ring systems. Building block 949 emerged as an enriched scaffold, appearing in 4 of the top 10 disynthons, consistently in combination with either a thiophene or an isoxazole (**Figure S8**). For HSA, we observe a majority being exclusively nitrogen bearing aromatic ring systems, with substituents of the rings being either primary amines, or methyl amides (**Figure S9**). Full features and documentation provided at https://github.com/SztainLab/DEL-iver/ include step-by-step tutorials for analyzing and visualizing DEL screening results, including individual building block and disynthon based hit-picking, visualization, and hit-expansion via predicting hits from make-on-demand libraries. We anticipate this will serve as a resource for analysis and assessing protein-ligand prediction methods that will continue to expand over time.

## Conclusion

Computational prediction of protein-ligand interactions continues to play an important role in modern drug discovery workflows, yet selecting appropriate modeling strategies for new screening campaigns remains challenging. In this work, we systematically evaluated ML and physics-based methods across DEL datasets to better characterize their strengths, limitations, and practical applicability. From this assessment we summarize three key takeaways: 1. Simple ML models can accurately predict binding of unseen molecules if they are chemically similar to the training distribution. For example, those composed of the same building blocks in unique combinations with the same DEL synthesis scheme. Predictions worsen further from the training distribution, such as unseen building blocks, and linear versus branched synthesis schemes. 2. DEL data is heavily imbalanced, with the majority of compounds being non-hits. Around 90% of the non-hits could be removed before significant performance drop was detected, making the most informative dataset much smaller than previously thought. 3. Using state-of-the-art docking and co-folding methods outperforms ligand-only ML models at predicting binding of OOD ligands in some cases, but performs near random in others. This is dependent on both the target and ligand chemical space. For example, Boltz-2 performed well at discriminating BRD4 binders in DEL compounds which did not contain a triazine core, while GaLigandDock was superior at discriminating sEH binders of this type. Adding docking features to the ML models offers modest improvement, but is limited by pose accuracy. To address the persistent challenge of generalizability, we advocate for rigorous pilot testing to define the boundaries and scope of each method for a given target and structural class prior to embarking on large-scale prediction campaigns. Our open-source DEL-iver package offers a guide to streamlining these efforts.

## Methods

### Data & Data Processing

Data was obtained from the NeurIPS 2024 - Predict New Medicines with BELKA page on the Kaggle website.^29^ RDKit (release 2025.09.03)^55^ substructure matching was used to classify molecules with and without a triazine scaffold. As an additional OOD set, an openly accessible DEL screen against sEH from Mobley and colleagues^27^ was used. This dataset was preprocessed by the authors and labeled with binary hit/non-hit values. Only BB SMILES (opposed to full molecule SMILES) including 10,010,000 non-hits and 116,666 hits were used in this study.

### Molecule Embeddings

RDKit was used to generate all of the following embeddings. Input molecules were encoded as either (1) whole-molecule ECFP4 fingerprints (1024 or 2048 bits, radius=2) or (2) concatenated building block fingerprints (BB1, BB2, BB3; each 1024 bits, radius=2), yielding a 3072-dimensional input vector for the latter case. Other chemical encodings including APDP,^56^ MACCS,^43^ and FCFP4^36^ were also tested alone or in combination with the ECFP4 fingerprints. Unless otherwise noted, all trained models described in the manuscript use BB ECFP4 fingerprint encodings. Embeddings can be calculated with the DEL-iver package via the *molecules* module and are automatically calculated when running *DEL_iver_results*.*py*.

### MLP Architectures

All models were implemented in PyTorch using the Adam optimizer (learning rate = 0.001) with either binary cross-entropy loss or binary cross-entropy with logit loss. Random forests and graph neural networks were considered, but performed significantly worse than MLPs, so the majority of models tested had MLP architectures. All MLPs consisted of an input and output layer, with between two and six hidden layers. Dropout with a probability of 0.5 and batch normalization were also considered, but marginally decreased performance, so were not used in the baseline model. Permutation invariance was applied via shared processing layers for BB2 and BB3 followed by an attention layer. The performance of the model before and after permutation invariance was applied can be found in (**Table S2**).

Using the hyperparameters and model architecture that performed best on the public test set, we selected the following core model: the model first processes BB1 through a dedicated linear layer, followed by ReLU activation. The input and output sizes of this layer are 1024. BB2 and BB3 are processed via a common linear layer followed by ReLU activation, sharing weights to ensure symmetry with respect to BB2 and BB3. The input and output sizes of this layer are 1024. An attention layer then aggregates the processed BB2 and BB3 features into a single 1024-dimensional representation. The output from the BB1 processing layer is then concatenated with this BB2/BB3 representation, resulting in a 2048-dimensional representation. This representation is then passed through a series of four fully connected hidden layers, decreasing the dimensions by half at each layer until a 256-dimensional representation is reached, at which point a linear layer outputs a 1-dimensional value. Every layer is followed by ReLU activation function, except for the output layer, which is followed by the sigmoid activation function. This model was trained using the binary cross-entropy with logit loss.

All models were trained with a batch size of 10000 for 10 epochs. For each epoch, the PyTorch dataloader randomly selected batches from the training data until all data was seen. Model training and hyperparameter scanning can be carried out with the DEL-iver python package via the *model* module. The baseline model used in the paper is the *BBFP_PermInvarNN_v3*, and is available in the DEL-iver package, but the default model is the *BBFP_NN_v1* model, as it does not assume permutation invariance is relevant for all libraries. Training this model is automatically performed when running the *del_iver_models*.*py* script.

### Data Splitting

In addition to training and test splits provided by the competition, several other split strategies were implemented.

#### Constructing non-hit down-sampled training sets

For each of the three protein targets, the different percentages of non-hit data were removed from the original training set. The corresponding test set was the same as the original test set. This process was repeated three times for each percentage removed to construct six sets of down-sampled training sets per percentage removed.

#### Random splitting of data

For random splitting, the non-hits in the original training set were down-sampled to include only 10% of all non-hits. The down-sampled training set was combined with the original test set, shuffled, and then randomly sampled to construct train and test sets (80/20 split) This process was repeated six times to construct six sets of random train/test sets.

#### Constructing non-triazine (OOD) spiked training sets

15,891 (∼10%) non-triazine molecules from the original test set were withheld for testing. Train sets were constructed to contain 10%, 25%, 50%, 75%, or 100% of the remaining non-triazine molecules. For each percentage, this process was repeated three times to construct three training sets per percentage of OOD molecules. These training subsets were then used to train two sets of MLP models. The first set of models was trained using only the OOD datasets, while the second set of models was trained using an aggregation of the OOD datasets added to the original training dataset.

Specification of different train/test splits can be carried out by the *dataloader* module in the DEL-iver package. A random 80/20 train/test split is used automatically when running the *DEL_iver_results*.*py* script.

### Identifying primary binding sites with DiffDock-L

To determine the primary binding site for sEH, an apo structure was folded from sequence with AlphaFold2 (AF2).^6^ This was done to prevent introduction of bias from crystallized structures with bound ligands. The sequence was retrieved from PDB entry 3WKE. DiffDock-L^57^ was then used to generate 100 binding pocket predictions for every ligand in the OOD set, using the default inference parameters provided in the DiffDock-L github repository. For each ligand, the DiffDock-L prediction with the highest internal DiffDock-L confidence score was compared to the known binding pockets for sEH. To compare to known binding pockets, the RMSD of the DiffDock-L prediction and the endogenous ligand in the known binding pockets were calculated with the PyMol CLI. The binding pocket with the smallest RMSD to the DiffDock-L prediction was used as the template binding pocket for docking the ligand with GALigandDock and Glide. Less than 1% of ligands bound to the secondary sEH pocket.

### Preparing docking template structures

Prior to all docking, the molecule SMILES strings were modified to replace the DNA attachment site on BB1 with a polyethylene glycol (PEG3) linker, i.e. replacing ‘[Dy]’ with ‘CCOCCOCCO’. This prevented modeling interactions with BB1 that would not be detected in the DEL screen. Though the exact linker composition and length used in the libraries is not known, PEG or is commonly used in DELs ^58^. Schrödinger Glide and Rossetta GALigandDock both require definition of a binding site. A single docked structure was used as a template for large-scale docking. For Glide, this controls the position of the docking grid centered around the ligand center of mass. For GALigandock, each new ligand is aligned with the ligand in the template structure using the PyMOL commandline interface to provide a starting conformation for docking perturbations. The template structure was generated using the ligand with the highest individual disynthon binding probability (Pbind) ^27^, determined for each disynthon by dividing the number times it is present in a hit by the number of times it is present in the entire dataset. For docking to sEH, compound 189754:

C#CC[C@@H](CC(=O)N[Dy])Nc1nc(NCC2CCC3(CCC3)CO2)nc(NCC23CC4CC(CC(C4)C2)C3)n1

was used with PDB: 3WKE.^59^ The compound was aligned in PyMOL to the co-crystalized ligand ‘AUB.’ The PEG3 linker was manually adjusted to point outward. Docking to BRD4 used PDB:8EWV,^60^ which contains a co-crystallized DEL molecule X5K for alignment. Only chain D in the PDB file was used for alignment and docking. Compound 91516204:

C#Cc1ccc(Nc2nc(Nc3cc(N4CCNCC4)ccc3[N+](=O)[O-])nc(N[C@@H](CC(=O)N[Dy])c3cccc(Cl)c3Cl)n2)cc1

was used as a template. Initial tests with Glide revealed the choice of initial template structure substantially impacts score distributions. First Glide Induced Fit Docking (IFD) was used to select a template for sEH. After generating 26 structures, two were selected for further analysis, 1) the template with the lowest docking score and 2) the template with the largest PEG SASA, indicating the PEG lies outside the pocket. Glide Extra Precision (XP) docking was then carried out with a random subset of the OOD set including 190 non-hits and all 175 hits. This revealed substantial differences in hit-discrimination capacity depending on the starting template (**Figure S10**). Glide Standard Precision (SP) docking was also tested and showed worse discrimination compared to Glide XP (**Figure S11**). Therefore, Glide XP was used for subsequent large-scale docking. To standardize the docking templates for large-scale docking, Rosetta GALigandDock^49,50^ with the dockflex method was used to generate templates as described in the next section. Following a single docking run where GALigandDock produced 20 poses per ligand, the molecule with the lowest energy score was chosen as the template. Selection was contingent on a plausible binding pose, ensuring the PEG linker tail remained oriented outside of the pocket rather than buried within it.

### Docking with Rosetta GALigandDock

Rosetta version 2020.08 was used. Ligands were prepared following the GALigandDock preprocessing guide available on RosettaCommons. First, ligand SMILES strings were converted to 3D conformations using Open Babel. Those that could not be generated with Open Babel were generated with RDKit. Next, Open Babel was used to add hydrogen atoms at pH 7.4. Partial charges were calculated with the Antechamber module of Amber24 using the AM1-BCC charge model. Parameters for Rosetta were generated with Rosetta’s mol2genparams.py script. Any molecule that produced an error during this preparation pipeline was excluded from the final docking set. This resulted in the removal of 2,021 sEH and 1,374 BRD4 molecules from the OOD set, alongside 776 sEH and 360 BRD4 from the ID set. Docking was performed with the RosettaScripts^31^ XML interface. The beta_genpot weights were used for docking scorefunction with the Lenard-Jones repulsive term weighted to 0.2. The beta_genpot_cart weights were used for the relax scorefunction. The GALigandDock protocol was then called with the dockflex method to allow for receptor flexibility. At each step of the docking, a genetic algorithm is employed in which the ligand is encoded into a gene with 6 rigid body degrees of freedom (DOFs) alongside additional DOFs representing internal bond rotations. The gene has a 20% chance of mutation and 80% of random crossover. For each conformation generated, a Monte Carlo algorithm search of receptor side chain rotameric space is carried out followed by minimization of both the receptor and the ligand. This generates 100 parent structures and 100 children structures, the 100 best scoring of these continue on as next the next iterations parents. After 10 iterations, the top 20 best scoring structures are side-chain optimized followed by backbone and side-chain minimization.^30^ The top ranked structure from these was used for all subsequent analysis.

### Docking with Schrödinger Glide

Schrödinger version 2024_02 was used. Protein structures from docking templates described above were prepared with the Schrödinger suite built-in tool Protein Preparation Wizard. This step involves adding missing hydrogens and protonation state assignment to the protein at the physiological pH of 7.4, and conducts restrained minimization to relax the protein and avoid clashes using OPLS-2005 force-field. The center of mass of the template molecules were used as the centroid for defining a cubic binding site. The Receptor Grid Generation module was then used to generate the docking grid while keeping the box size similar to the molecule in the template structure.

A cubic inner box of 10x10x10 by default was specified to constrain the ligand center during the exhaustive search for both sEH and BRD4. A cubic outer box was used to define the volume for grid potential calculations, with dimensions of approximately 29 Å for BRD4, 33 Å for the sEH hydrolase site, and 28 Å for the sEH phosphatase site. Molecules were prepared with the LigPrep module, which generates 3D structures from SMILES, adds hydrogens, and minimizes energy. Epik was used to predict the protonation states of molecules, and up to 32 stereoisomers were generated for each input structure. All the 3D conformers were retained for extensive binding pose sampling. Docking was carried out using the Glide module with Extra Precision (XP) mode. Only the docking pose with the best docking score for each molecule was used for the analysis.

### Co-folding with Boltz-2

Boltz-2^8^ was used to co-fold ligands in the OOD test set, and 25K ligands from the ID set to each of the three protein targets. As was done for docking, the DNA tag in each molecule was replaced by a PEG3 linker prior to Boltz-2 inference. Default parameters for running inference with Boltz-2 were used, and no constraints on binding pockets were used. The binding affinity and the binding probability were predicted with each of the co-folded structures. The MSAs for each protein were precalculated and then provided for each subsequent co-folding job to increase the speed of jobs. For BRD4, UniProt IDs M0QZD9 (residues 59-164) was used for generating MSAs and predicted structures. For sEH, the fasta from PDB 3WKE was used. After finishing the calculations, an extra three residues were found on the N-terminal domain (QTN) as a result of a pasting error, as these are the first three residues of BRD4. These terminal residues are on the secondary domain, not the active site domain, therefore this should not impact the results. Given that the PLIPs only consider residues which interact with the ligand, this should not impact those either.

### Docking analysis

To interpret the docking scores from different molecular docking methods as ligand-binding probabilities, raw scores were sign-inverted and linearly normalized to the [0, 1] interval. Under this transformation, better (more negative) docking scores correspond to higher probabilities. Other methods were tested to define binding probability including assigning a score cutoff for binary labeling, and only performing linear scaling on percentage of scores above a threshold. None of these performed better, and thus were not used. Binding poses were compared across docking methods by calculating the ligand RMSD after alignment of protein Cα atoms CPPTRAJ ^61^ (**Figure S4**). Solvent accessible surface area (SASA) shown in **Figure 4** was calculated using the rdFreeSASA RDKit module. Prior to calculation, ProDy^62^ was used to convert PDB files to RDKit molecules.

### Protein-Ligand Interaction (PLIP) fingerprints of docked structures

PLIP analysis was performed for each docked protein structure with the plip python package,^53^ using the default parameters. Fingerprints were then computed with the output PLIP xml files by determining the residue contacts over all ligands in a docked set, then creating a bit vector of the length of interacting residues, where 1 represents ligand interaction at the corresponding residue, and 0 represents no ligand interaction at the corresponding residue. For each protein target, the fingerprints were computed across all ligands bound to the target so that PLIP fingerprints would have a consistent length. To integrate structural features into ML model training, the PLIP fingerprints were concatenated with ECPF4 fingerprints. The baseline ML architecture was modified to accommodate the larger input layer size. Similar to the baseline, the input was passed through a series of four fully connected hidden layers, decreasing the dimensions by half at each layer.

## Data Availability

Data from the BELKA competition is available at: https://polarishub.io/datasets/leash-bio/belka-v1. Docked and co-folded structures generated in this study are available at: https://huggingface.co/datasets/SztainLab/BELKA_docked_structures

## Code Availability

DEL-iver code is available at: https://github.com/SztainLab/DEL-iver. Schrödinger Glide is available at: https://www.schrodinger.com/. Rosetta is available at: https://rosettacommons.org/software/. Boltz-2 is available at: https://github.com/jwohlwend/boltz

## Acknowledgements

We thank Leash Biosciences for hosting the BELKA Kaggle competition and making their data available for this study. This material is based upon work supported by the U.S. Department of Energy, Office of Science, Office of Advanced Scientific Computing Research, Department of Energy Computational Science Graduate Fellowship under Award Number DE-SC0025528 to M.D and National Institute of Health R35 GM151129 to M.J.O.

## Author Contributions

T.S. conceptualized the study and supervised the project. M.D., D.S.P, Y.F, SH.L., and S.M. developed code and conducted computational experiments. All authors contributed to data analysis and manuscript preparation.

## Notes

### Competing Interest Statement

The authors have declared no competing interest.

### Summary of Updates

Modified some language to clarify the broad applications to protein-ligand binding prediction from high throughput screening in general, though specifically applied here to the unique challenges associated with DNA-encoded library screens.

