## Supplementary Information for "Assessing the Generalizability of Machine Learning and Physics-Based Methods with DNA-Encoded Libraries"

### Table of Contents

1. **Supplementary Figures**

**Figure S1.** Performance of model on additional OOD dataset.
**Figure S2.** Protein ligand interaction profiles (PLIPs) interaction counts from docked structures.
**Figure S3.** Performance of structure-based methods in the ID BEKLA set and analysis of and training on physics-derived features.
**Figure S4.** Pairwise RMSD between predicted poses from each physics-based method.
**Figure S5**. Distribution of building blocks and their corresponding pBind values for each target.
**Figure S6.** Disynthon pBind for BRD4, sEH, and HSA.
**Figure S7.** Top 10 building blocks and disynthons for sEH ranked by pBind from the BELKA dataset.
**Figure S8.** Top 10 building blocks and disynthons for BRD4 ranked by pBind from the BELKA dataset.
**Figure S9.** Top 10 building blocks and disynthons for HSA ranked by pBind from the BELKA dataset.
**Figure S10.** Impact of the docking template on hit discrimination.
**Figure S11.** Comparison of Glide SP and Glide XP on hit discrimination.

2. **Supplementary Tables**

**Table S1.** Results of ML architecture design decisions.
**Table S2.** Enrichment factors for physics-based methods on the OOD set.
**Table S3.** Statistics for disynthon 2_3 distribution for molecules with more than 30 occurrences.

#### Supplementary Figures

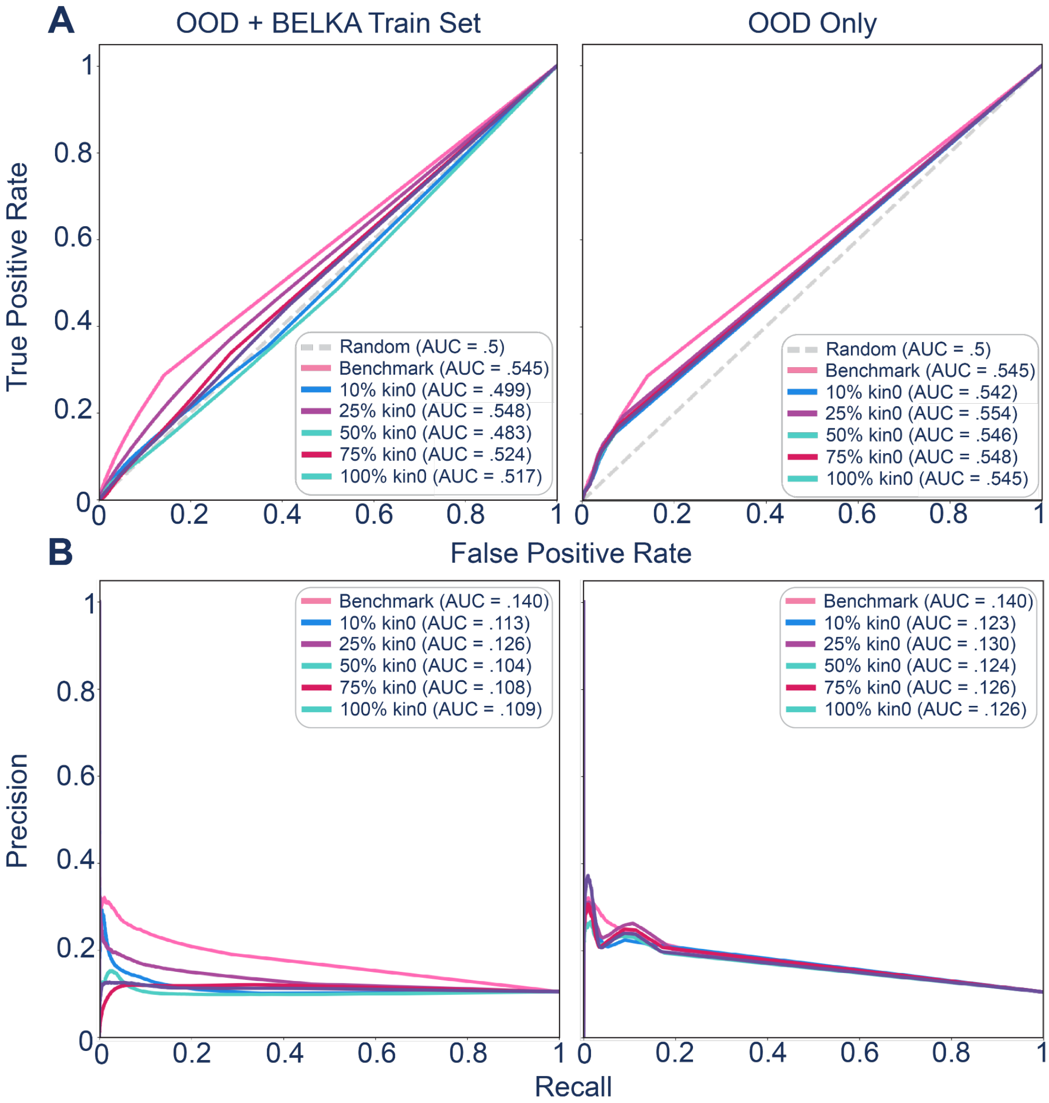


**Figure S1.** Performance of model on the additional OOD (Mobley) dataset. **A)** AUROC plots showing the performance of ML models on the Mobley dataset. The benchmark model is the fixed MLP model trained on the original BELKA train set. The additional models are trained on either the kin0 set aggregated to the original training dataset or on the kin0 set alone. **B)** The corresponding precision-recall curves to the AUROC plots in (**A**).

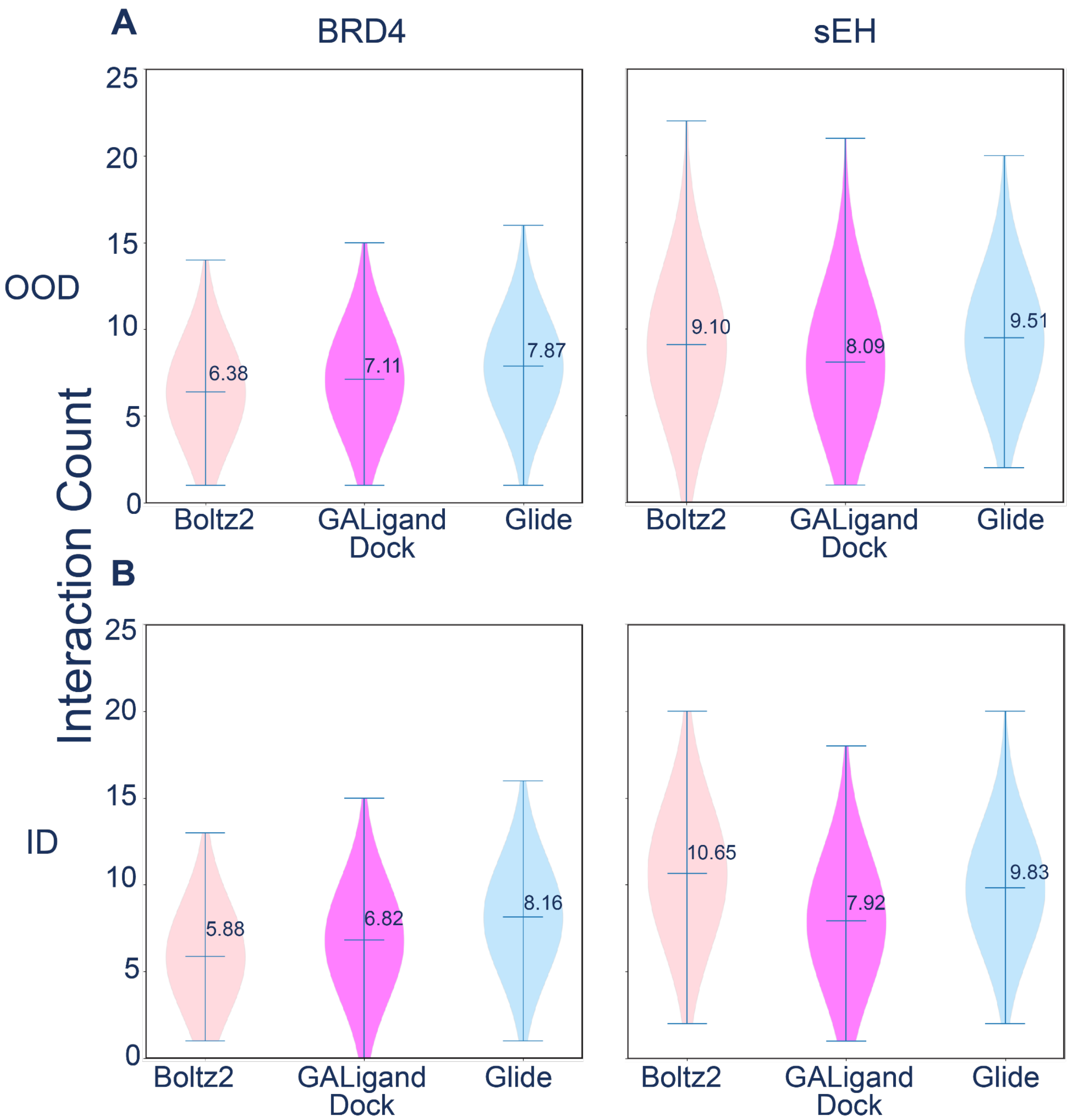


**Figure S2.** Protein ligand interaction profiles (PLIPs) interaction counts from docked structures. **A)** Interaction counts from PLIP fingerprints of OOD molecules docked to BRD4 (left) and sEH (right). **B)** Interaction counts from PLIP fingerprints of ID molecules docked to BRD4 (left) and sEH (right).

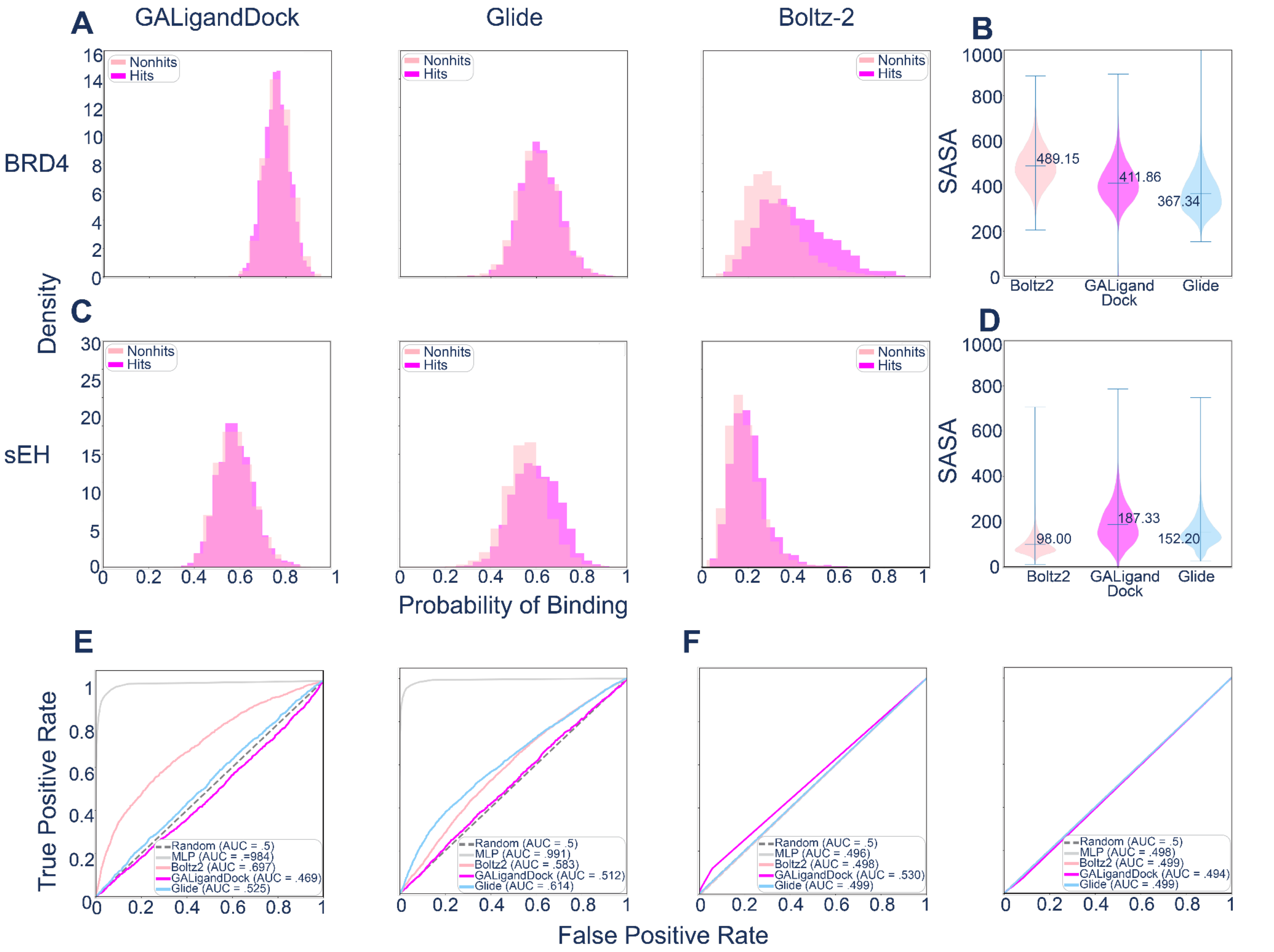


**Figure S3.** Performance of structure-based methods on the ID BELKA set and analysis of and training on physics-derived features. **A)** Probability of binding derived from GALigandDock, Glide, and Boltz-2 for the ID BRD4 set. **B)** SASA of the docked structures from GALigandDock, Glide, and Boltz-2 for the OOD BRD4 set. **C)** Probability of binding derived from GALigandDock, Glide, and Boltz-2 for the ID sEH set. **D)** SASA of the docked structures from GALigandDock, Glide, and Boltz-2 for the OOD sEH set. **E)** AUROC plots showing the performance of the baseline ML predictions and physics-based methods on predicting hits and non-hits for the BRD4 (left) and sEH (right) ID sets. **F)** AUROC plots showing the performance of the baseline ML architecture and ML architectures using concatenated PLIP and ECFP4 fingerprints as input. Models were trained on the 25k ID set and tested on the OOD set (BRD4: left, sEH: right). For each protein, the PLIPs calculated from each structure-based method were used to train a separate model.

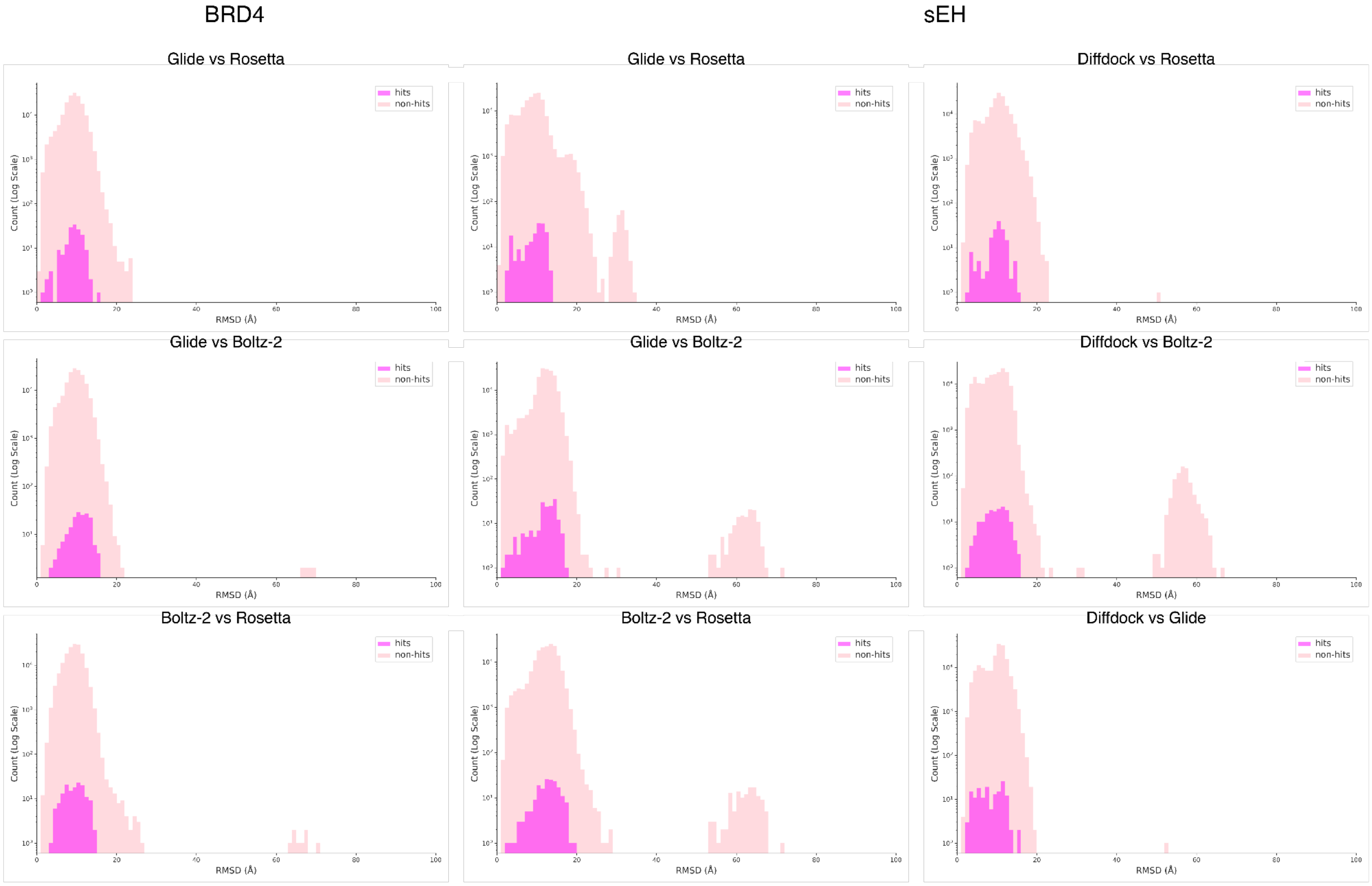


**Figure S4**. Pairwise RMSD between predicted poses from each method. For each pair, the receptor of one predicted complex was aligned to the other based on Cα atoms. The heavy-atom RMSD of the ligands was then calculated using CPPTRAJ.

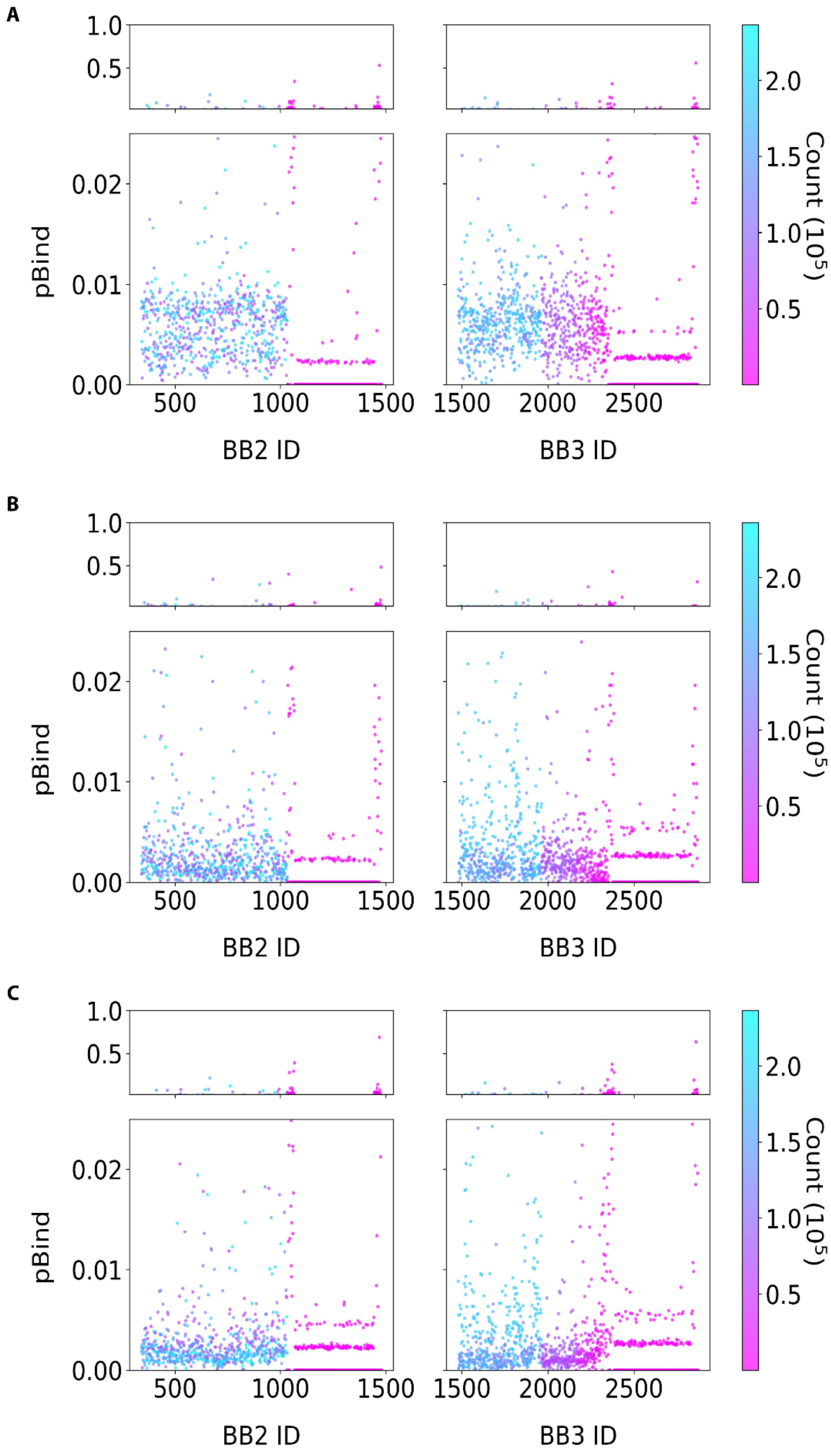


**Figure S5**. Distribution of building blocks and their corresponding pBind values for each target. (A) sEH, (B) BRD4, and (C) HSA.

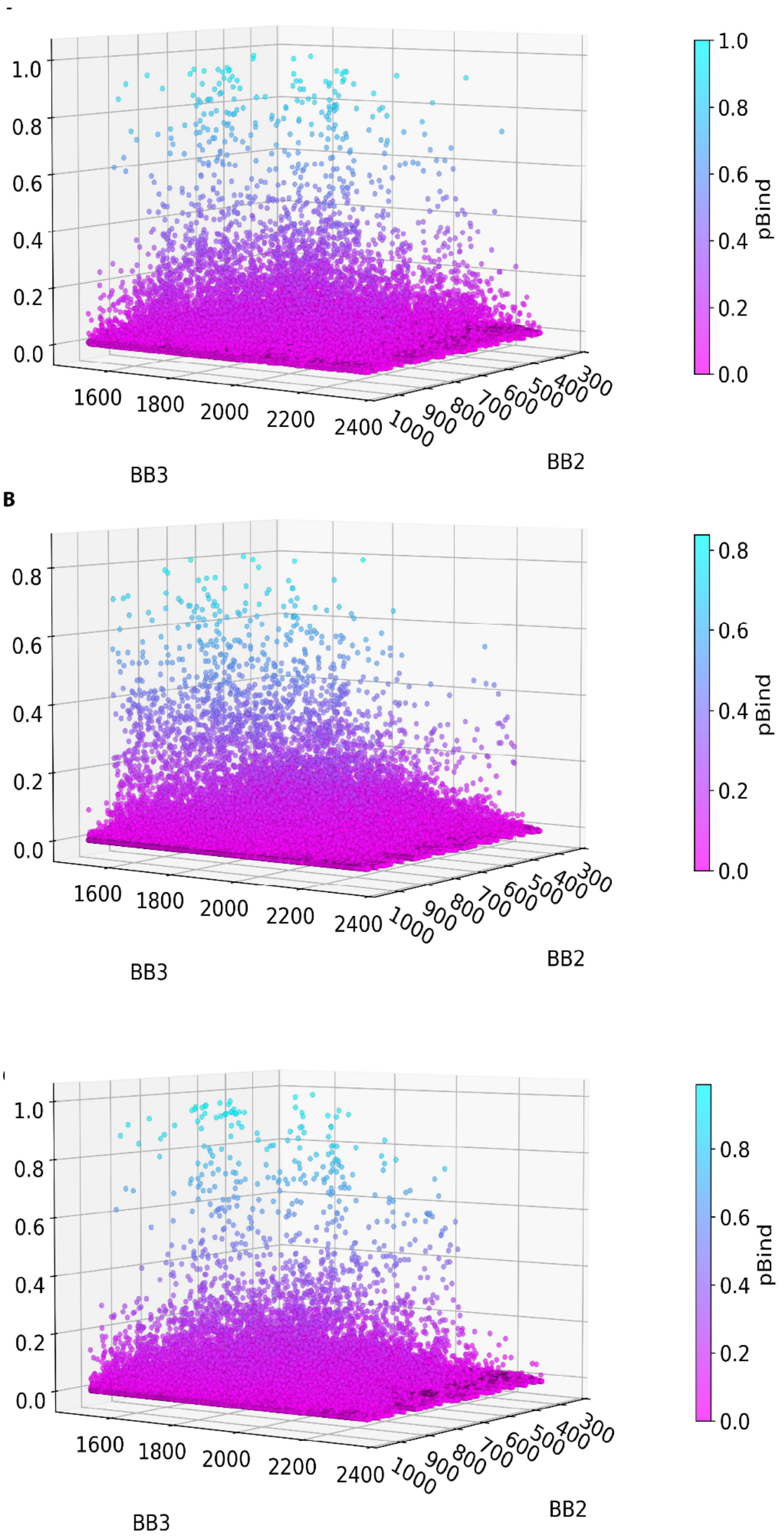


**Figure S6**. Distribution of disynthons and their corresponding pBind values for each target. (A) sEH, (B) BRD4, and (C) HSA.

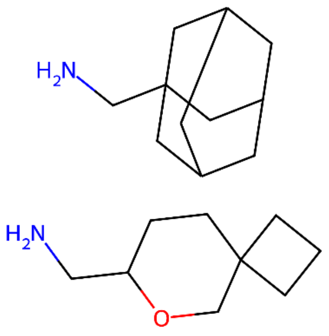

ID: 189754 | pbind: 1.00
BB2(704) + BB3(696)

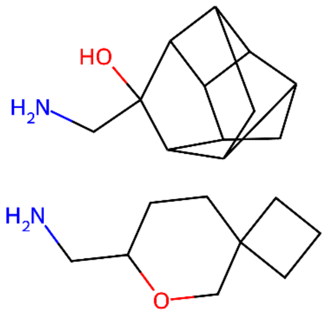

ID: 189752 | pbind: 0.99
BB2(704) + BB3(694)

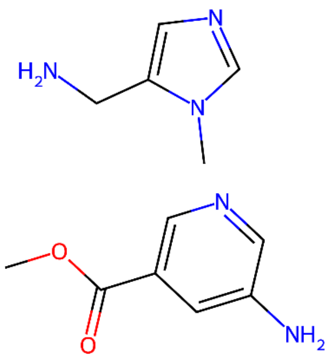

ID: 65293 | pbind: 0.98
BB2(464) + BB3(634)

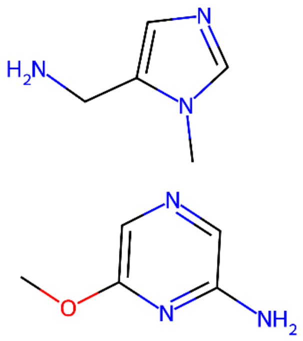

ID: 99134 | pbind: 0.97
BB2(528) + BB3(634)

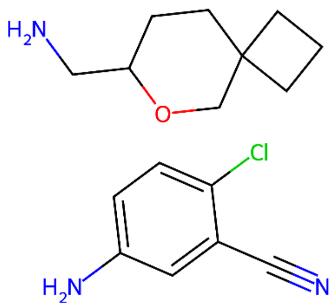

ID: 235305 | pbind: 0.97
BB2(795) + BB3(704)

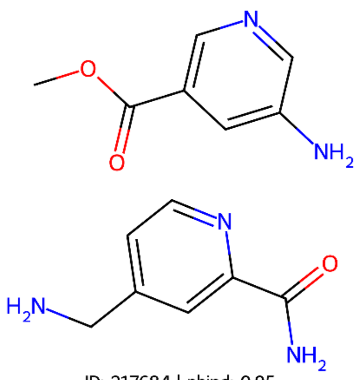

ID: 217684 | pbind: 0.95
BB2(759) + BB3(464)

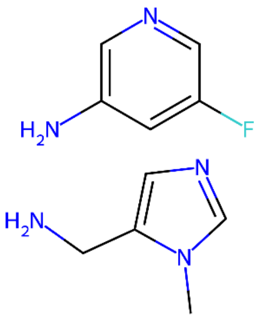

ID: 152939 | pbind: 0.95
BB2(634) + BB3(994)

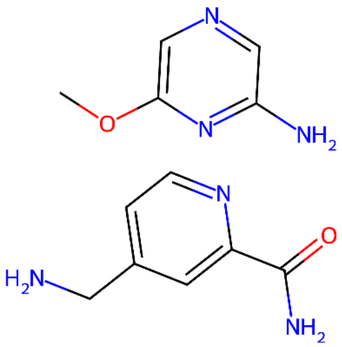

ID: 217750 | pbind: 0.94
BB2(759) + BB3(528)

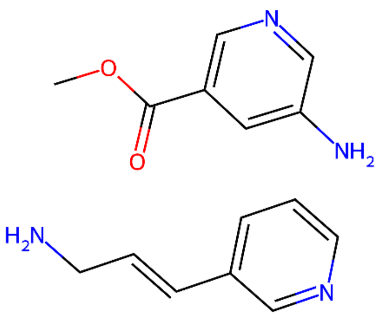

ID: 157650 | pbind: 0.94
BB2(643) + BB3(464)

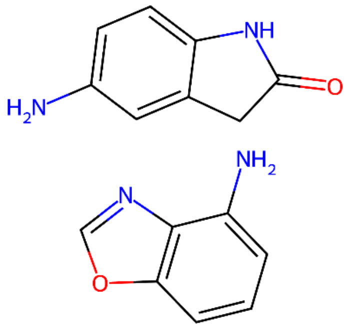

ID: 333391 | pbind: 0.94
BB2(975) + BB3(946)

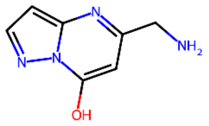

ID: 1470 | Position: BB3, BB2 | pbind: 0.56, 0.53

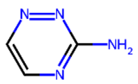

ID: 1067 | Position: BB2, BB3 | pbind: 0.34, 0.32

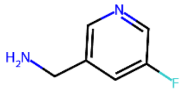

ID: 666 | Position: BB2, BB3 | pbind: 0.19, 0.13

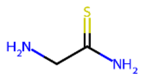

ID: 1462 | Position: BB2 | pbind: 0.16

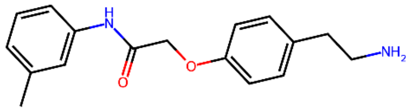

ID: 1044 | Position: BB3, BB2 | pbind: 0.16, 0.11

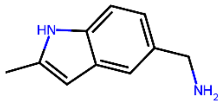

ID: 1532 | Position: BB3 | pbind: 0.15

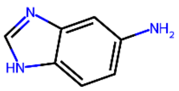

ID: 1063 | Position: BB3, BB2 | pbind: 0.15, 0.12

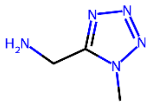

ID: 1459 | Position: BB3 | pbind: 0.15

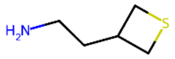

ID: 834 | Position: BB2, BB3 | pbind: 0.11, 0.11

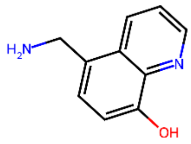

ID: 659 | Position: BB2 | pbind: 0.11

**Figure S7**. Top 10 building blocks and disynthons for sEH ranked by pBind from the BELKA dataset. Structures were generated from SMILES using the MolFromSmiles and MolsToGridImage functions in RDkit.

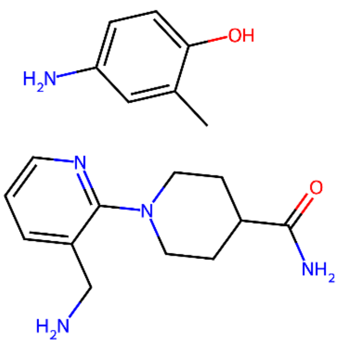

ID: 276333 | pbind: 0.84
BB2(867) + BB3(573)

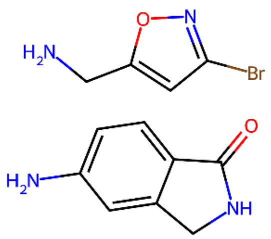

ID: 319878 | pbind: 0.83
BB2(949) + BB3(1560)

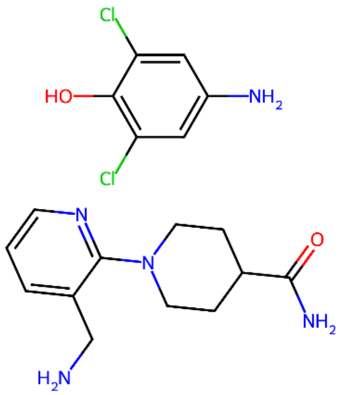

ID: 276659 | pbind: 0.83
BB2(867) + BB3(886)

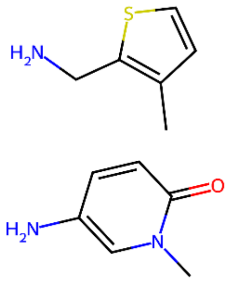

ID: 177808 | pbind: 0.82
BB2(680) + BB3(1540)

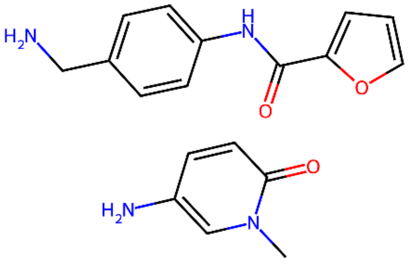

ID: 177884 | pbind: 0.81
BB2(680) + BB3(1597)

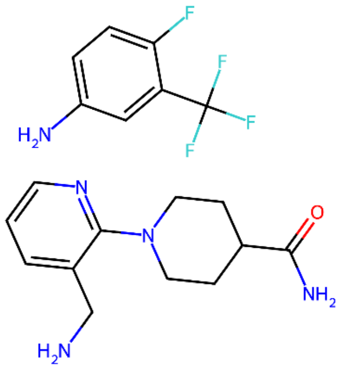

ID: 276688 | pbind: 0.80
BB2(867) + BB3(918)

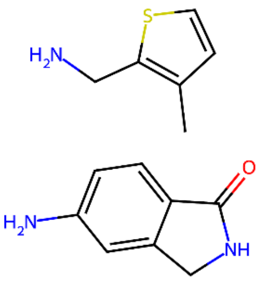

ID: 319852 | pbind: 0.79
BB2(949) + BB3(1540)

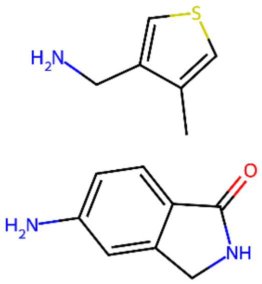

ID: 319857 | pbind: 0.79
BB2(949) + BB3(1545)

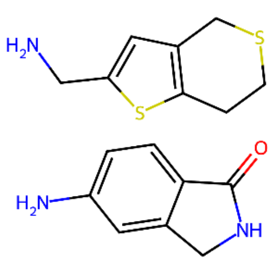

ID: 319922 | pbind: 0.78
BB2(949) + BB3(1592)

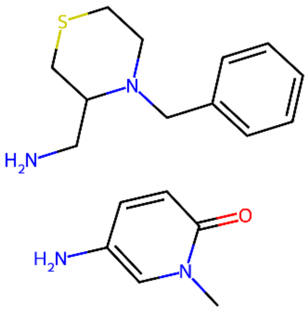

ID: 177869 | pbind: 0.75
BB2(680) + BB3(1584)

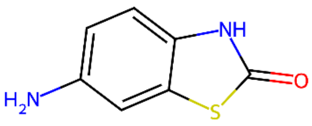

ID: 1478 | Position: BB2, BB3 | pbind: 0.48, 0.31

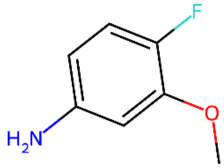

ID: 1039 | Position: BB3, BB2 | pbind: 0.43, 0.40

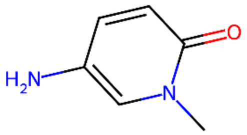

ID: 680 | Position: BB2, BB3 | pbind: 0.34, 0.19

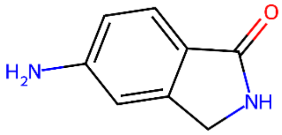

ID: 949 | Position: BB2, BB3 | pbind: 0.29, 0.09

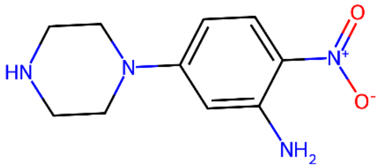

ID: 901 | Position: BB2, BB3 | pbind: 0.27, 0.25

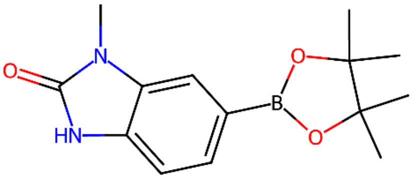

ID: 1337 | Position: BB2 | pbind: 0.22

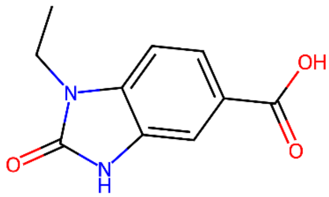

ID: 1706 | Position: BB3 | pbind: 0.13

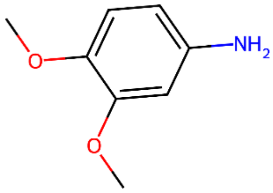

ID: 508 | Position: BB2 | pbind: 0.11

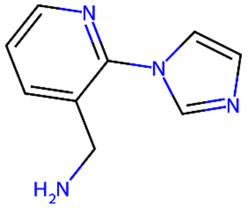

ID: 1475 | Position: BB2 | pbind: 0.10

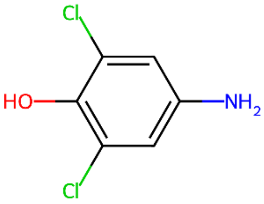

ID: 886 | Position: BB2 | pbind: 0.09

**Figure S8**. Top 10 building blocks and disynthons for BRD4 ranked by pBind from the BELKA dataset. Structures were generated from SMILES using the MolFromSmiles and MolsToGridImage functions in RDkit.

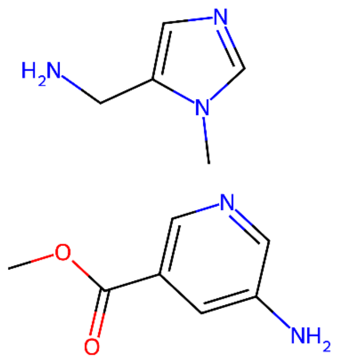

ID: 65293 | pbind: 0.99
BB2(464) + BB3(634)

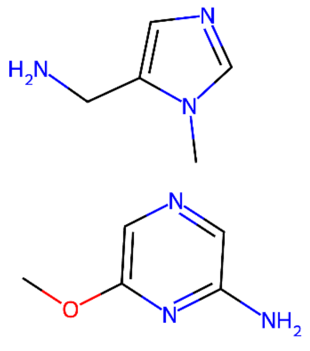

ID: 99134 | pbind: 0.99
BB2(528) + BB3(634)

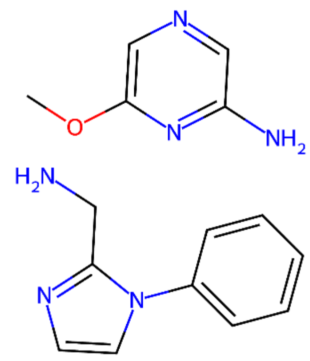

ID: 174525 | pbind: 0.98
BB2(674) + BB3(528)

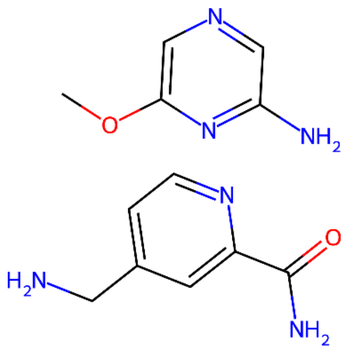

ID: 217750 | pbind: 0.97
BB2(759) + BB3(528)

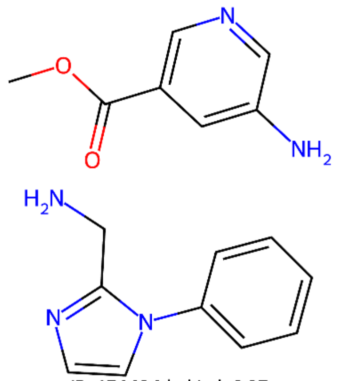

ID: 174494 | pbind: 0.97
BB2(674) + BB3(464)

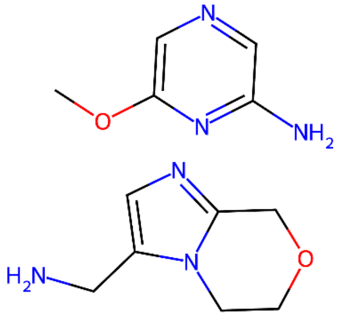

ID: 218882 | pbind: 0.97
BB2(762) + BB3(528)

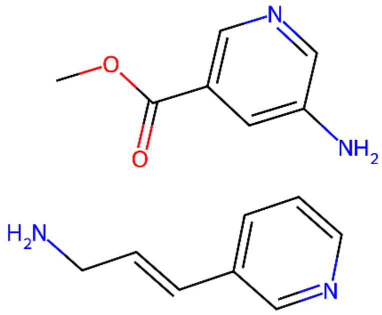

ID: 157650 | pbind: 0.96
BB2(643) + BB3(464)

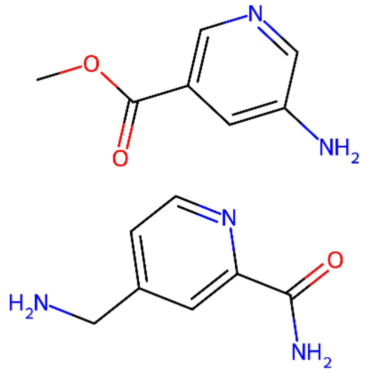

ID: 217684 | pbind: 0.96
BB2(759) + BB3(464)

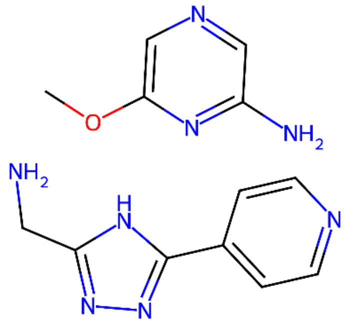

ID: 223829 | pbind: 0.96
BB2(772) + BB3(528)

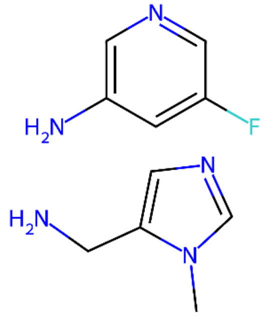

ID: 152939 | pbind: 0.96
BB2(634) + BB3(994)

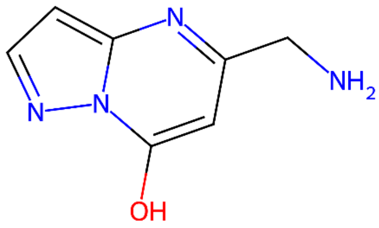

ID: 1470 | Position: BB2, BB3 | pbind: 0.69, 0.64

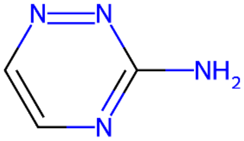

ID: 1067 | Position: BB2, BB3 | pbind: 0.39, 0.37

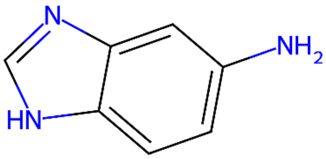

ID: 1063 | Position: BB3, BB2 | pbind: 0.31, 0.29

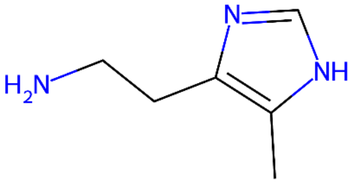

ID: 1043 | Position: BB3, BB2 | pbind: 0.28, 0.28

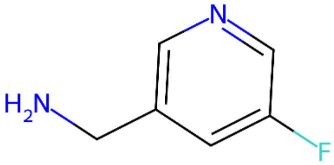

ID: 666 | Position: BB2, BB3 | pbind: 0.22, 0.16

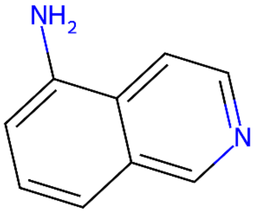

ID: 973 | Position: BB3 | pbind: 0.20

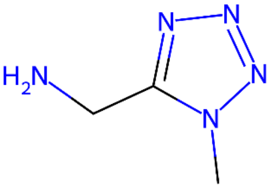

ID: 1459 | Position: BB3 | pbind: 0.19

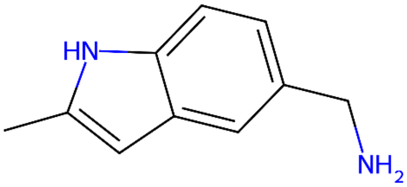

ID: 1532 | Position: BB3 | pbind: 0.16

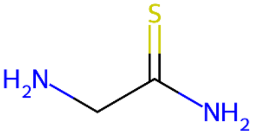

ID: 1462 | Position: BB2 | pbind: 0.14

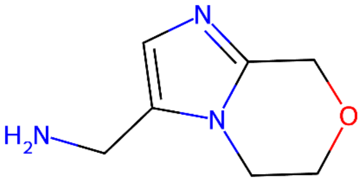

ID: 762 | Position: BB3 | pbind: 0.13

**Figure S9**. Top 10 building blocks and disynthons for HSA ranked by pBind from the BELKA dataset. Structures were generated from SMILES using the MolFromSmiles and MolsToGridImage functions in RDkit.

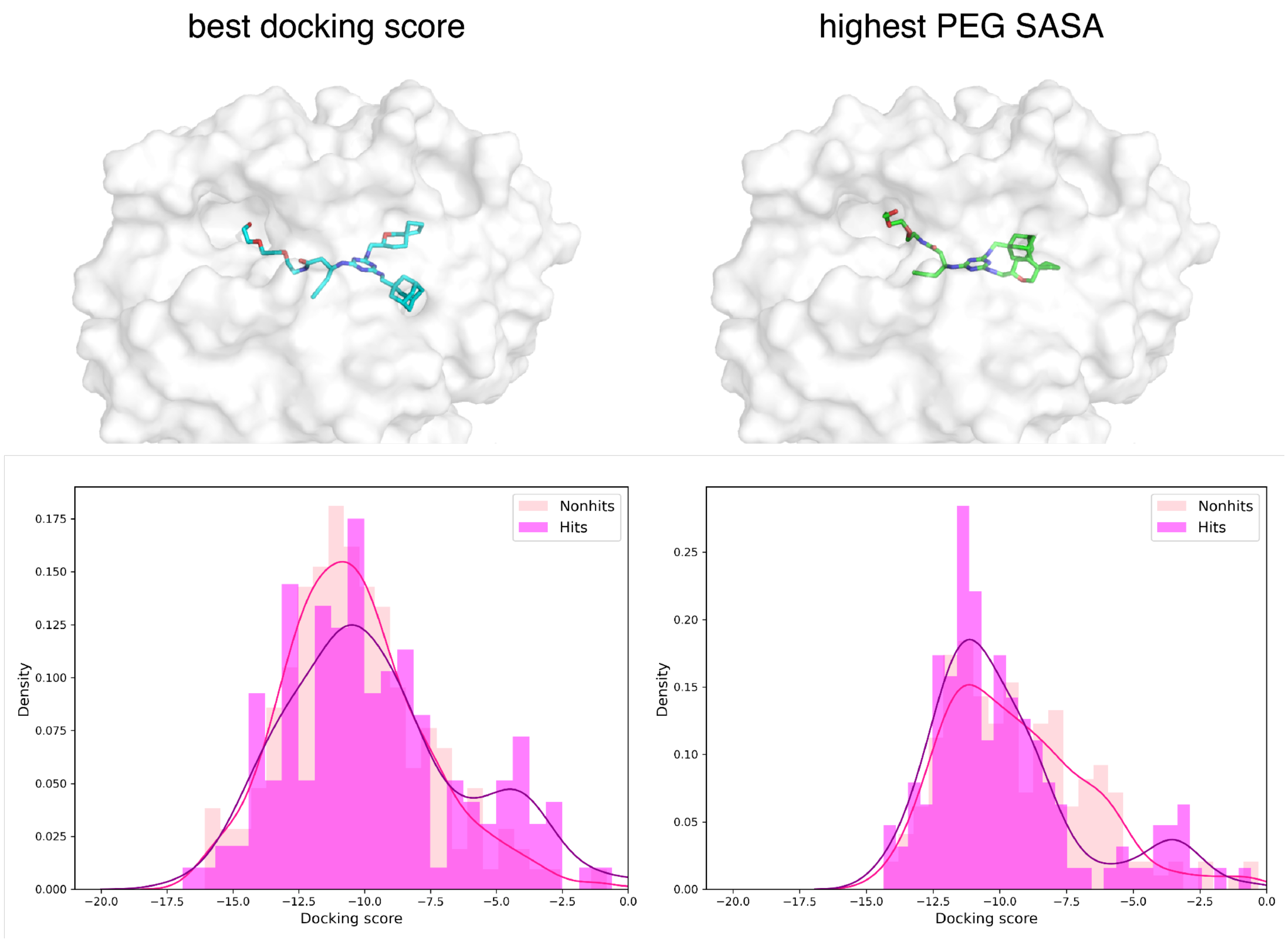


**Figure S10**. Impact of docking template on hit discrimination. Top: Comparison of two template structures generated with Glide IFD. A single ligand with the highest pBind was docked to sEH resulting in 26 output structures. The structure with the best docking score, and the structure with the largest PEG linker SASA were chosen as templates for docking 190 non-hits and all 175 hits from the OOD set. Bottom: Score distribution of hits and non-hits from Glide XP docking to each template.

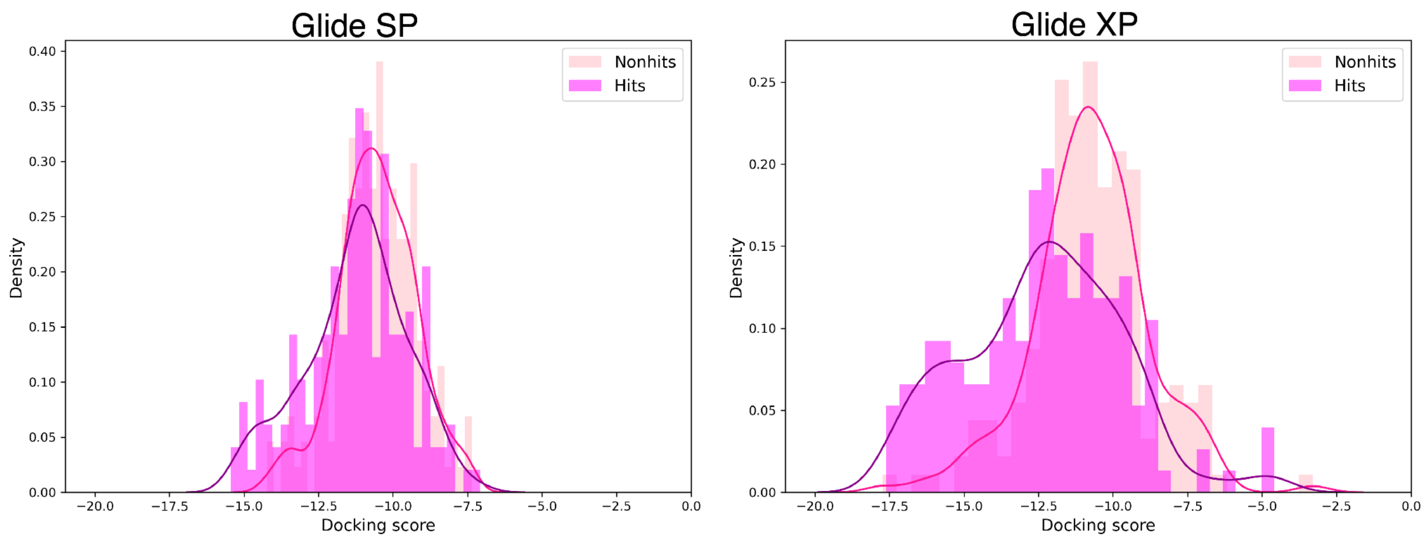


**Figure S11.** Comparison of Glide SP and Glide XP hit discrimination. The two methods were compared using the same compound set in Figure S10, using an identical docking grid for consistency. In contrast to S10, the template used for these tests was generated using GALigandDock for clearer comparison between the two methods.

#### Supplementary Tables

**Table S1.** Results of ML architecture design decisions. All models listed here were trained with 10 epochs, learning rate 0.001, and batch size 10000. The starred model is the baseline model used throughout the paper. The last two rows introduce permutation invariance in two distinct ways. The first concatenates BB2 and BB3 fingerprints before input into the model, while the second passes each ECFP4 fingerprint separately, but processes them through a shared layer.

| Model | Hidden layers | mAP public | mAP private |
|---|---|---|---|
| **Initial MLP w/ ECFP4s** | [512, 256] | 0.321 | 0.163 |
| **MACCS FPs** | [256, 128] | 0.335 | 0.220 |
| **FCFP4s** | [512, 256] | 0.332 | 0.213 |
| **Add dropout to ECFP4 model** | [1024, 512, 256] | 0.336 | 0.222 |
| **APDPs** | [2048, 1024, 512, 256] | 0.319 | 0.211 |
| **Concat APDPs and ECFP4s** | [2048, 1024, 512, 256] | 0.440 | 0.251 |
| **Concat BB2 and BB3 in input** | [512, 256] | 0.400 | 0.222 |
| **Shared BB2/BB3 processing layer + attention*** | [2048, 1024, 512, 256] | 0.434 | 0.243 |

**Table S2.** Enrichment factors for physics-based methods on the OOD. Enrichment factors are calculated from predicted binding probabilities from Boltz-2, GALigandDock, and Glide on the OOD and ID sets for BRD4 and sEH.

| Set | EF0.5% | EF1% | EF5% | EF10% | EF20% |
|---|---|---|---|---|---|
| **BRD4 OOD Boltz-2** | 86.04 | 55.38 | 16.34 | 9.08 | 9.08 |
| **BRD4 OOD GALigandDock** | 0 | 0.54 | 0.97 | 0.91 | 0.91 |
| **BRD4 OOD Glide** | 4.30 | 3.23 | 1.51 | 1.34 | 1.34 |
| **sEH OOD Boltz-2** | 6.86 | 6.86 | 4.34 | 2.74 | 2.74 |
| **sEH OOD GALigandDock** | 9.15 | 10.28 | 8.34 | 6.63 | 6.63 |
| **sEH OOD Glide** | 9.15 | 8.0 | 4.80 | 3.43 | 3.43 |
| **BRD4 ID Boltz-2** | 4.37 | 3.68 | 3.77 | 3.11 | 3.11 |
| **BRD4 ID GALigandDock** | 0.59 | 0.81 | 0.92 | 0.94 | 0.94 |
| **BRD4 ID Glide** | 1.23 | 1.08 | 1.11 | 1.15 | 1.15 |
| **sEH ID Boltz-2** | 1.36 | 1.32 | 1.41 | 1.33 | 1.33 |
| **sEH ID GALigandDock** | 1.38 | 1.38 | 1.02 | 0.98 | 0.98 |
| **sEH ID Glide** | 1.84 | 1.76 | 2.16 | 2.08 | 2.08 |

**Table S3**. Statistical overview of disynthons derived from BB2 and BB3, limited to those occurring more than 30 times.

| Metric | Mean | Median | 90th% | 99th% | Max | Protein |
|---|---|---|---|---|---|---|
| **pBind** | 0.005 | 0 | 0.007 | 0.092 | 0.838 | BRD4 |
| **N hits** | 1.258 | 0 | 2.0 | 25.0 | 227 | BRD4 |
| **Ntotal** | 271.0 | 271.0 | 271.0 | 271.0 | 271.0 | BRD4 |
| **pBind** | 0.007 | 0.004 | 0.007 | 0.085 | 1.0 | sEH |
| **N hits** | 1.997 | 1.0 | 2.0 | 23.0 | 271.0 | sEH |
| **Ntotal** | 271.0 | 271.0 | 271.0 | 271.0 | 271.0 | sEH |
| **pBind** | 0.004 | 0.0 | 0.007 | 0.085 | 0.989 | HSA |
| **N hits** | 1.124 | 0.0 | 2.0 | 23.0 | 268.0 | HSA |
| **Ntotal** | 271.0 | 271.0 | 271.0 | 271.0 | 271.0 | HSA |
